# SARS-CoV-2 ORF8 sequence conservation and mutational analysis — insight into the influence of dataset size on identifying top mutations

**DOI:** 10.1101/2025.06.16.659836

**Authors:** Shubhangi Kandwal, Ivan Cmelo, Darren Fayne

## Abstract

Given how quickly the SARS-CoV-2 virus mutates, the COVID-19 pandemic has been a major source of concern. The ORF8 accessory protein is one such protein, which is reported to have undergone many mutations. This makes ORF8 an intriguing protein to investigate how these mutations might play a role in overall ORF8 activity. In this study, we have performed conservation and mutational analysis on SARS-CoV- 2 ORF8 protein sequences to identify the conserved and mutated residues. We have also split the ORF8 sequence data into SARS-CoV-2 variant datasets to further identify top mutations across each of them. The mutated and conserved residues were visualised on the available structure of ORF8 to highlight the conserved and mutated sites, which might hold some biological significance. Finally, our study also investigated the significance of sequence dataset size in capturing top mutations following multiple sequence alignments.

**Author Summary:** The COVID-19 pandemic was caused by the SARS-CoV-2 virus, which is known to change over time, i.e., it gets mutated, resulting in the generation of different variants. The ORF8 accessory protein of the SARS-CoV- 2 genome is known to undergo these changes more frequently. In our study, we used SARS-CoV-2 ORF8 protein sequences from various variants to identify mutations among them. Furthermore, we have discovered sites that remain unchanged over time, a phenomenon known as conservation. We think that these unchanged and changed sites could be important for biology and studying them will help in understanding the underlying mechanism of how ORF8 interacts with partner proteins based on existing experimental data. Lastly, we have looked at how much sequence data is sufficient for identifying the top mutated sites.

**Graphical abstract:** 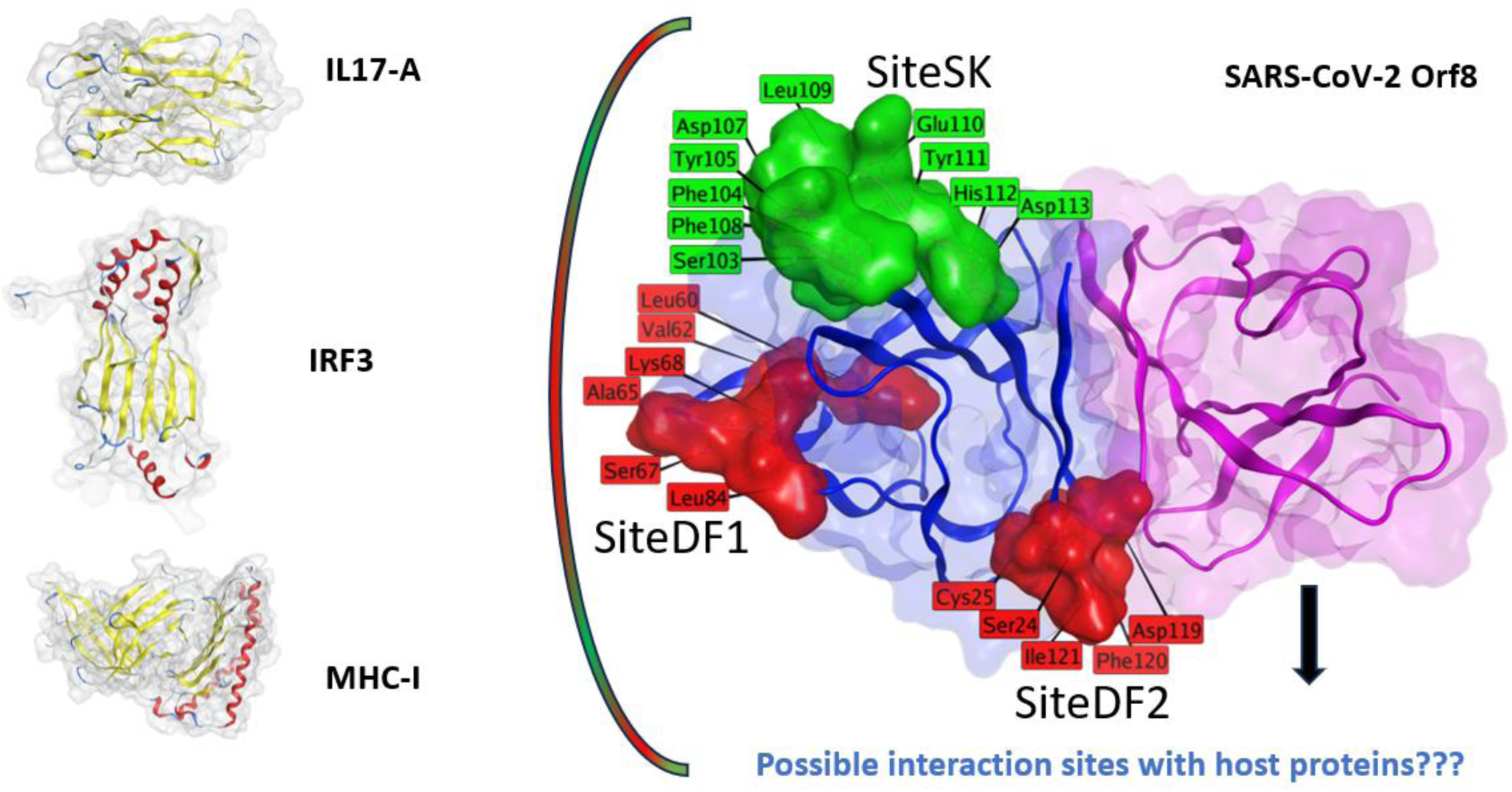

## 1. Introduction

The severe acute respiratory syndrome coronavirus 2 (SARS-CoV-2) (SC-2) is a Betacoronavirus belonging to the Coronaviridae family and caused the coronavirus disease 2019 (COVID-19) pandemic. The SC-2 genome is a large positive-sense single-stranded RNA that encodes for 29 proteins, including four structural, sixteen non-structural, and nine accessory proteins (Zhou et al., 2020). The non-structural proteins (NSPs) are considerably more conserved across coronaviruses (CoVs) (Naqvi et al., 2020), in particular, pairwise sequence analysis between the NSPs of SARS-CoV-1 (SC-1) and SC-2 demonstrated that there is a high level of sequence conservation between both viruses (Kandwal & Fayne, 2023) and they could represent viable targets for drug repurposing studies (Kandwal & Fayne, 2022). For example, proteins such as NSP3 and NSP13 are known to be highly conserved not only across SC-2 sequences but also across other betacoronaviruses such as SC-1 and the Middle East respiratory syndrome coronavirus (MERS). Interestingly, the CACHE consortium selected both of these NSPs as targets in the second and third rounds of a recently initiated international hit-finding competition (https://cache-challenge.org/)(Herasymenko et al., 2025).

The ORF8 gene is present within a cluster of accessory genes such as ORF6, ORF7a and ORF7b which is located just after the M gene and preceding the N gene (Figure 1). Among the accessory proteins, ORF8 has 366 nucleotides (positions 27,894-28,259), which encode for 121 amino acids (Supplementary Material Excel sheet S01). The ORF8 gene is poorly conserved and has been associated with SC-2 virulence (Pereira, 2020).

**Figure 1.**
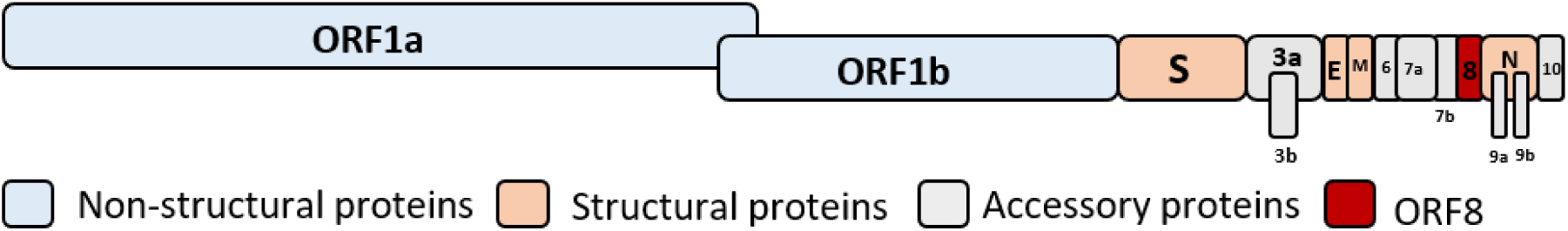
SARS-CoV-2 genome showing different coding regions (structural, non-structural, and accessory proteins).

ORF8 has an unstructured N-terminal signal sequence with an Ig-like fold (Tan et al., 2020). The ORF8 protein is comprised of an N-terminal hydrophobic signal peptide (1-15 AA) and an ORF8 chain (16-121 AA) (Hassan et al., 2021). The SC-2 ORF8 protein has a novel ability to dimerise compared to other Sarbecoviruses. The monomeric units are comprised of an α-helix, followed by a β-sandwich of seven β-strands, which are stabilized through three intramolecular disulfide bonds (C25-C90, C37-C102, C61-C83) (X. Chen et al., 2021) (Valcarcel et al., 2021) (Flower et al., 2021). The dimerization of the two monomers is also facilitated by intermolecular disulfide bonding between C20 residues, stabilizing the dimer interface (Flower et al., 2021).

The specific role of ORF8 is still not fully understood, but a few studies have indicated possible cellular roles. Upon SC-2 infection, ORF8 helps SC-2 to evade the host immune system by downregulating the expression of major histocompatibility complex class 1 (MHC-I) (Zhang et al., 2021) along with the suppression of type 1 interferon antiviral response (Rashid, Dzakah, et al., 2021). A few findings have also suggested that ORF8 interacts with IL-17 to induce IL-17RA-mediated inflammation (Lin et al., 2021) (Wu et al., 2022). In another study, SC-2 ORF8 was reported to inhibit type 1 interferon (IFN-β) activation and the NF-κB pathway, thereby affecting the host antiviral response (Li et al., 2020). The ORF8 protein modulates the unfolded protein response (UPR) by up-regulating ER-resident chaperones GRP78 and GRP94, thus stimulating the ATF6 and IRE1 pathways (Valcarcel et al., 2021).

The protein sequence alignment suggests that SC-2 ORF8 shares homology with Bat-RaTG13-CoV ORF8, as it has a high sequence similarity (95%) but, surprisingly, a low similarity with SC-1 ORF8 (30%) (Lu et al., 2020). The first comparison between the genomic regions of SC-1 and SC-2 placed ORF8 among the most divergent genes (Wu et al., 2020). It is well known that viruses change over time, leading to the emergence of variants having different viral properties, such as how they spread, the severity of disease caused upon infection, or response to therapeutic agents. The World Health Organization (WHO) assigned labels such as variants of concern (VOCs) and variants of interest (VOIs) representing different levels of severity, and the major VOCs discussed in this paper are Alpha, Beta, Gamma, Delta and Omicron.

The ORF8 gene is reported to be a hypervariable region undergoing rapid nucleotide substitutions and deletions (S. Chen et al., 2020; Pereira, 2020). This hypervariability of ORF8 is exemplified across many studies. An Alpha variant having 382-nucleotide deletions (Δ382) in the ORF7b and ORF8 genes (Su et al., 2020) resulted in no expression of ORF8, which was then associated with mild COVID-19 symptoms in patients (Young et al., 2020). A few of the bioinformatics studies are summarized in Table 1, but, in brief, a recent study focused on reduced ORF8 expression in SC-2 lineages due to mutations present in the transcriptional regulatory sequences (TRS) (Hisner et al., 2023) that caused truncations in the VOC Alpha ORF8 protein. An SC-2 ORF8 gene Multiple Sequence Alignment (MSA) study also showed the deletion of 382 nucleotides in the sequences extracted from two of the main sequence repositories: the National Centre for Biotechnology Information (NCBI) and the Global Initiative on Sharing All Influenza Data (GISAID) (Khare et al., 2021) (Mohammad et al., 2020).

**Table 1:**
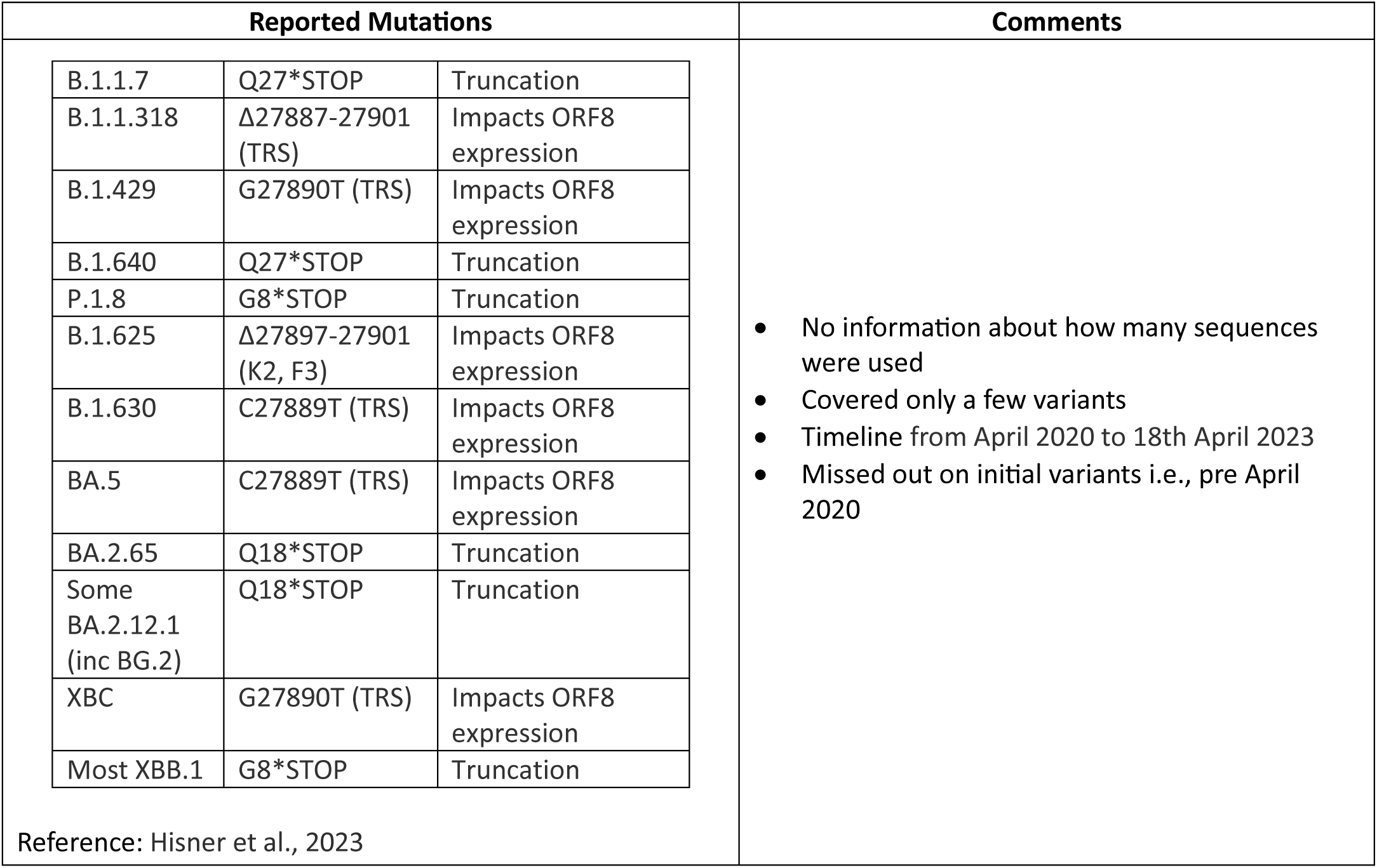
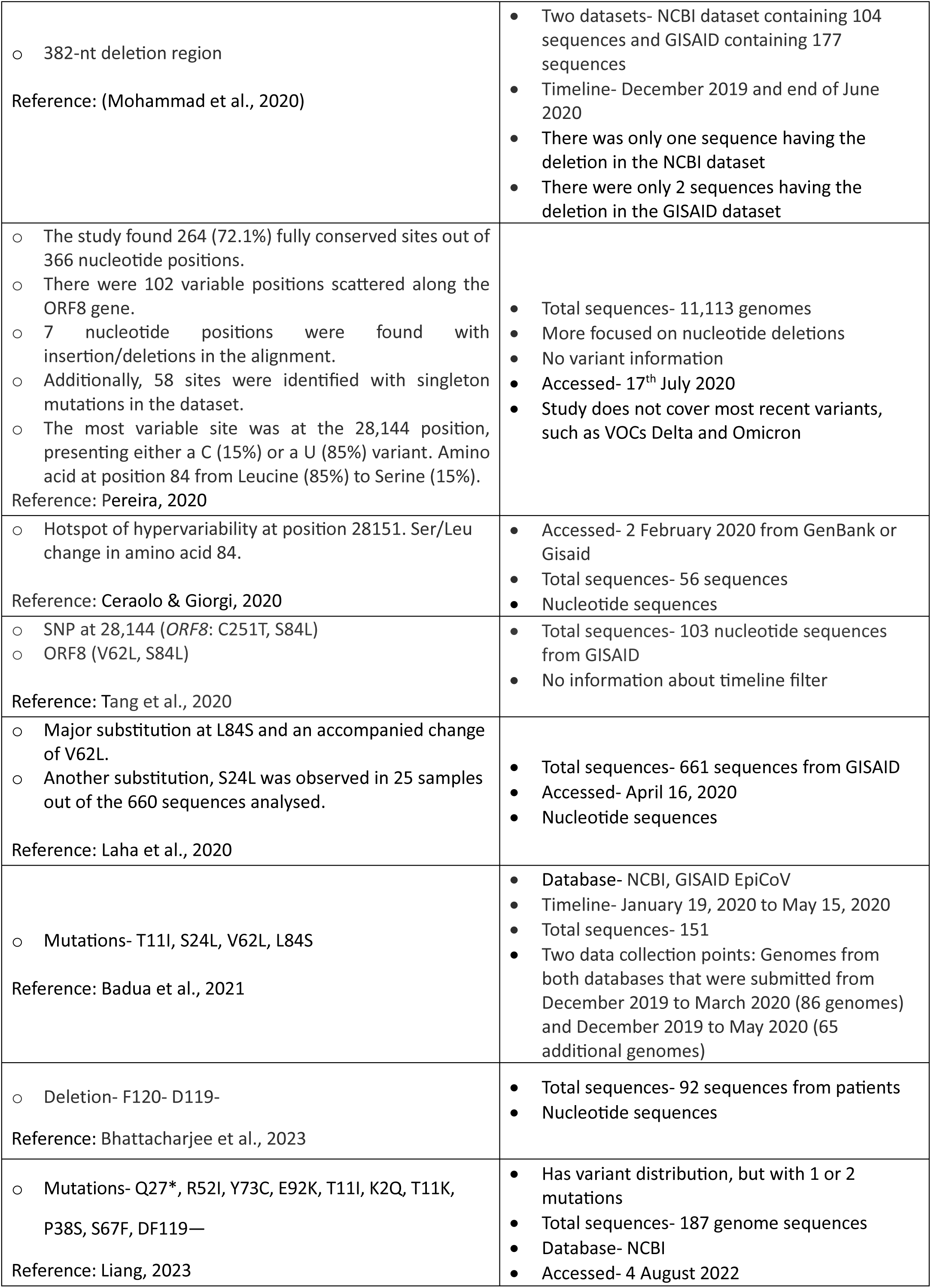
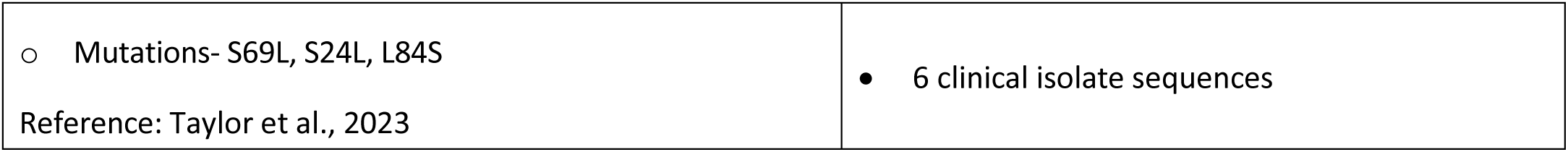
List of various SC-2 ORF8 studies.

| Reported Mutations |  |  | Comments |
| --- | --- | --- | --- |
| B.1.1.7 | Q27*STOP | Truncation | No information about how many sequences were used<br>Covered only a few variants<br>Timeline from April 2020 to 18th April 2023<br>Missed out on initial variants i.e., pre April 2020 |
| B.1.1.318 | Δ27887-27901 (TRS) | Impacts ORF8 expression |  |
| B.1.429 | G27890T (TRS) | Impacts ORF8 expression |  |
| B.1.640 | Q27*STOP | Truncation |  |
| P.1.8 | G8*STOP | Truncation |  |
| B.1.625 | Δ27897-27901 (K2, F3) | Impacts ORF8 expression |  |
| B.1.630 | C27889T (TRS) | Impacts ORF8 expression |  |
| BA.5 | C27889T (TRS) | Impacts ORF8 expression |  |
| BA.2.65 | Q18*STOP | Truncation |  |
| Some BA.2.12.1 (inc BG.2) | Q18*STOP | Truncation |  |
| XBC | G27890T (TRS) | Impacts ORF8 expression |  |
| Most XBB.1 | G8*STOP | Truncation |  |
| Reference: Hisner et al., 2023 |  |  |  |

Table 1: List of various SC-2 ORF8 studies.
|  |  |
| --- | --- |
| ○ 382-nt deletion region<br>Reference: (Mohammad et al., 2020) | • Two datasets- NCBI dataset containing 104 sequences and GISAID containing 177 sequences<br>• Timeline- December 2019 and end of June 2020<br>• There was only one sequence having the deletion in the NCBI dataset<br>• There were only 2 sequences having the deletion in the GISAID dataset |
| ○ The study found 264 (72.1%) fully conserved sites out of 366 nucleotide positions.<br>○ There were 102 variable positions scattered along the ORF8 gene.<br>○ 7 nucleotide positions were found with insertion/deletions in the alignment.<br>○ Additionally, 58 sites were identified with singleton mutations in the dataset.<br>○ The most variable site was at the 28,144 position, presenting either a C (15%) or a U (85%) variant. Amino acid at position 84 from Leucine (85%) to Serine (15%).<br>Reference: Pereira, 2020 | • Total sequences- 11,113 genomes<br>• More focused on nucleotide deletions<br>• No variant information<br>• Accessed- 17<sup>th</sup> July 2020<br>• Study does not cover most recent variants, such as VOCs Delta and Omicron |
| ○ Hotspot of hypervariability at position 28151. Ser/Leu change in amino acid 84.<br>Reference: Ceraolo & Giorgi, 2020 | • Accessed- 2 February 2020 from GenBank or GisaId<br>• Total sequences- 56 sequences<br>• Nucleotide sequences |
| ○ SNP at 28,144 (*ORF8*: C251T, S84L)<br>○ *ORF8* (V62L, S84L)<br>Reference: Tang et al., 2020 | • Total sequences- 103 nucleotide sequences from GISAID<br>• No information about timeline filter |
| ○ Major substitution at L84S and an accompanied change of V62L.<br>○ Another substitution, S24L was observed in 25 samples out of the 660 sequences analysed.<br>Reference: Laha et al., 2020 | • Total sequences- 661 sequences from GISAID<br>• Accessed- April 16, 2020<br>• Nucleotide sequences |
| ○ Mutations- T11I, S24L, V62L, L84S<br>Reference: Badua et al., 2021 | • Database- NCBI, GISAID EpiCoV<br>• Timeline- January 19, 2020 to May 15, 2020<br>• Total sequences- 151<br>• Two data collection points: Genomes from both databases that were submitted from December 2019 to March 2020 (86 genomes) and December 2019 to May 2020 (65 additional genomes) |
| ○ Deletion- F120- D119-<br>Reference: Bhattacharjee et al., 2023 | • Total sequences- 92 sequences from patients<br>• Nucleotide sequences |
| ○ Mutations- Q27*, R52I, Y73C, E92K, T11I, K2Q, T11K, P38S, S67F, DF119—<br>Reference: Liang, 2023 | • Has variant distribution, but with 1 or 2 mutations<br>• Total sequences- 187 genome sequences<br>• Database- NCBI<br>• Accessed- 4 August 2022 |
| ○ Mutations- S69L, S24L, L84S<br>Reference: Taylor et al., 2023 | • 6 clinical isolate sequences |

Expanding on the studies mentioned in Table 1, which clearly shows that most studies have identified mutations at position 84, with two major lineages identified: L84 and S84. Among the two lineages, the frequently emerged L84S ORF8 mutation leads to reduced IL-17RA binding on monocytes and attenuated inflammation in vitro (Wu et al., 2022). A study focused on aligning 11,113 ORF8 nucleotide sequences and found that the 28,144 site was the most variable site with the presence of either a C (15%) or a U (85%) (Pereira, 2020) which resulted in a change from leucine (in 85% of the sequences) to serine (in 15% of the sequences) at position 84. Another study found that the amino acid region between 83-89 is more likely to be disordered in circulating variants having serine at position 84 (Ceraolo & Giorgi, 2020) as it induces structural disorder in the C-terminal of the protein, which was determined by the Russell/Linding algorithm (Linding et al., 2003). A study by (Tang et al., 2020) also reported the presence of a nonsynonymous mutation at the same 28,144 genomic location (*ORF8*: C251T) encoding for L84 in 101 of 103 sequences, whereas only 29 sequences encoded for S84. The presence of these two amino acids at position 84 should be explored further to understand their functional importance. An *in-silico* based study performed a one microsecond molecular dynamics (MD) simulation to observe the dimeric behaviour of the L84S mutation in comparison to the wild-type protein and found that the L84S mutation affects overall structural stability by reducing the frequency of protein-protein interactions (PPIs) between the dimer (Islam et al., 2024). Interestingly, a few mutations might not occur independently but rather be accompanied by a corresponding substitution, for example, V62L with L84S (Laha et al., 2020), as noted following an alignment of 661 sequences. An MSA analysis of two full-length SC-2 sequence datasets (containing 86 and 65 sequences, respectively) at different time points across various geographical locations found that ORF8 presented the highest mutation density across the 29 SC-2 proteins. The study suggested the distinctive abundance of ORF8 mutations was quite similar in the two datasets (Badua et al., 2021). The ORF8 protein is also reported to have two deletions, which were identified at the end of the ORF8 protein, i.e., D119- and F120- (Bhattacharjee et al., 2023) in a 187-sequence MSA study. An interesting study reported that the Delta variant sequences had two deletions at DF119- - residue positions and the Alpha variant had truncated ORF8 proteins due to a stop codon at Q27* (Liang, 2023). A study by (Taylor et al., 2023) reported nonsynonymous mutations such as S69L, S24L and L84S associated with six clinical isolates.

All of these previous studies focused on small sequence datasets and did not explore the presence of particular mutations across all the different VOCs (Alpha, Beta, Gamma, Delta and Omicron) as well as non-VOCs. Usually, bioinformatics-based studies use an arbitrary selection of thresholds to classify amino acids/nucleotides as being mutated or conserved. For example, (Ou et al., 2022) identified novel mutations in the spike protein of Omicron subvariants if that particular mutation had a frequency of occurrence in > 50 sequences from a total of 52,563 sequences. Another study used 2% as the cutoff value for the registration of mutations from a dataset having 383,570 SC-2 genome sequences (Weber et al., 2021). A study by (Perlinska et al., 2022) also used an arbitrary selection of a mutation rate of > 0.002 for identifying frequent mutations from a dataset having 2,610,999 SC-2 PLpro sequences.

In our study, we have first identified mutations considering an arbitrary selection of mutation threshold (0.74%) from the MSA of over 1 million SC-2 ORF8 variant sequence datasets having varying amino acid lengths referred to as ORF8 (1M-varying AA) in this study in order to determine the top 10 mutations, followed by identifying mutations associated with different VOC datasets. Our dataset contained over one million sequences, which gave us the statistical confidence to identify the most populated mutations that could not be identified in smaller sequence datasets. The top 10 ORF8 mutations were identified across the variants, and the presence of each mutation was compared with other variant datasets. We have also identified highly conserved amino acid residues in the different datasets that fall within the arbitrarily selected conserved threshold value of 0.03%.

Most previous studies used small dataset sizes; indeed, small datasets are widely used across the bioinformatics literature, but we explored the importance of dataset size in MSA studies to identify the most prevalent mutations. In order to gain more insight into the question, “How large is large enough?”, we ran MSA studies (x100) on dataset sizes from 5 to 773,689, containing SC-2 ORF8 VOCs and non-VOCs sequences having 121 amino acid lengths, also referred to as ORF8 (700k-121 AA) sequences, and determined how large a dataset had to be sampled in order to accurately determine the most prevalent mutated AA positions. Although the original dataset contained over one million sequences, the MSA was negatively affected by the variability in sequence lengths, as there were large gaps. To compensate for this, the MSA output file was fine-tuned to identify mutations; however, the combination of varied lengths and random resampling in our second study introduced excessive gaps, ultimately compromising alignment quality and mutation detection. To mitigate these issues, we selected a high-quality subset of equal-length sequences, ensuring more consistent alignment and minimising artefacts. Our findings have wider implications and statistical analyses for consideration, in MSA studies focused on determining amino acid, or nucleotide, mutation rates.

## 2. Results

**(A) Conservation and mutational analysis**

### 2.1 ORF8 (1M-varying AA) sequence dataset MSA analysis

After pre-processing of the dataset removed 77 ambiguous sequences, the MSA of 1,010,757 ORF8 protein sequences resulted in a wide range of conservation scores at each amino acid position. The highest was 99.99% at the start M1 position, and the lowest was 76.55% at F120 position. The dataset had 773,689 sequences of 121 amino acid length, i.e., the length of the reference sequence (Wuhan strain) followed by 233,329 sequences of 119 amino acid length. The majority of the sequences in the dataset belong to the Omicron variant, followed by Delta, Non-VOC, Gamma, Beta and Alpha (Figure 2a). The top two deletions were F120- and D119- followed by eight missense mutations in order of prevalence, were S24L, T11I, L60F, E92K, Y31F, C25Y, A51S and A65V (Figure 2b).

**Figure 2.**
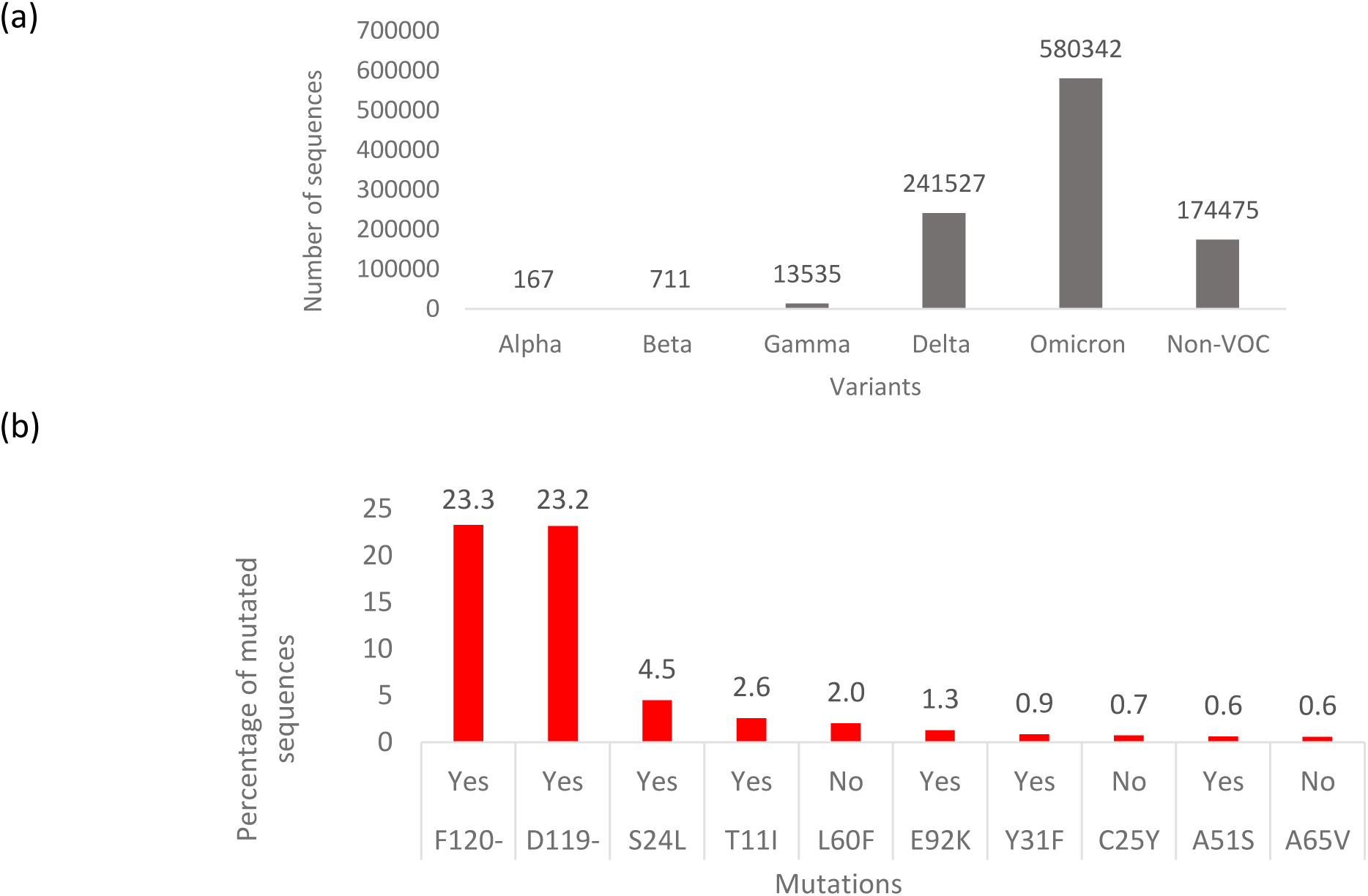
(a) Distribution of sequences across the variants. Alpha-B.1.1.7 and subvariants, Beta-B.1.352 and subvariants, Gamma-P.1, P.2 and subvariants, Delta-B.1.617.2 and subvariants, Omicron-BA.1 and subvariants and Non-VOCs. (b) Top 10 mutations or deletions from the ORF8 (1M-varying AA) alignment showing the percentage of sequences having the mutations in the protein dataset. Yes indicates a change in amino acid properties, No indicates no change in amino acid properties.

### 2.2 MSA analysis of VOCs

After analysing the full SC-2 ORF8 dataset, we created subdatasets of sequences belonging to particular variants and distributed sequences into two major categories, VOCs and non-VOCs. The VOCs comprised of those known before to July 30, 2022; Alpha, Beta, Gamma, Delta and Omicron. The splitting of the ORF8 (1M-varying AA) sequence dataset into the VOCs and non-VOCS resulted in varying numbers of sequences within each dataset, thus changing the mutation and conservation thresholds. So, for the variant analysis, we extracted the top 10 mutations occurring in the MSA of each variant dataset without considering the thresholds. To predict the possible impact of mutations on the protein, I-Mutant scores were calculated to estimate the structural stability, and the mutations were also visualized on 3D-protein structures. After determining the top 10 mutations for a particular variant, we analysed the presence of these mutations across the other variant datasets.

**(a) Alpha**

The Alpha variant dataset had a total of 102 sequences of length 121 AAs and 65 sequences with only 26 AAs (Figure 3a). The MSA was performed on the 102 sequences, and the only seven mutations found were V62L, Y73C, R52I, T11I, A65V, P70T and D119E (Figure 3b, 3c). I-Mutant scores suggested that the mutations decreased the stability of the protein structure.

**Figure 3.**
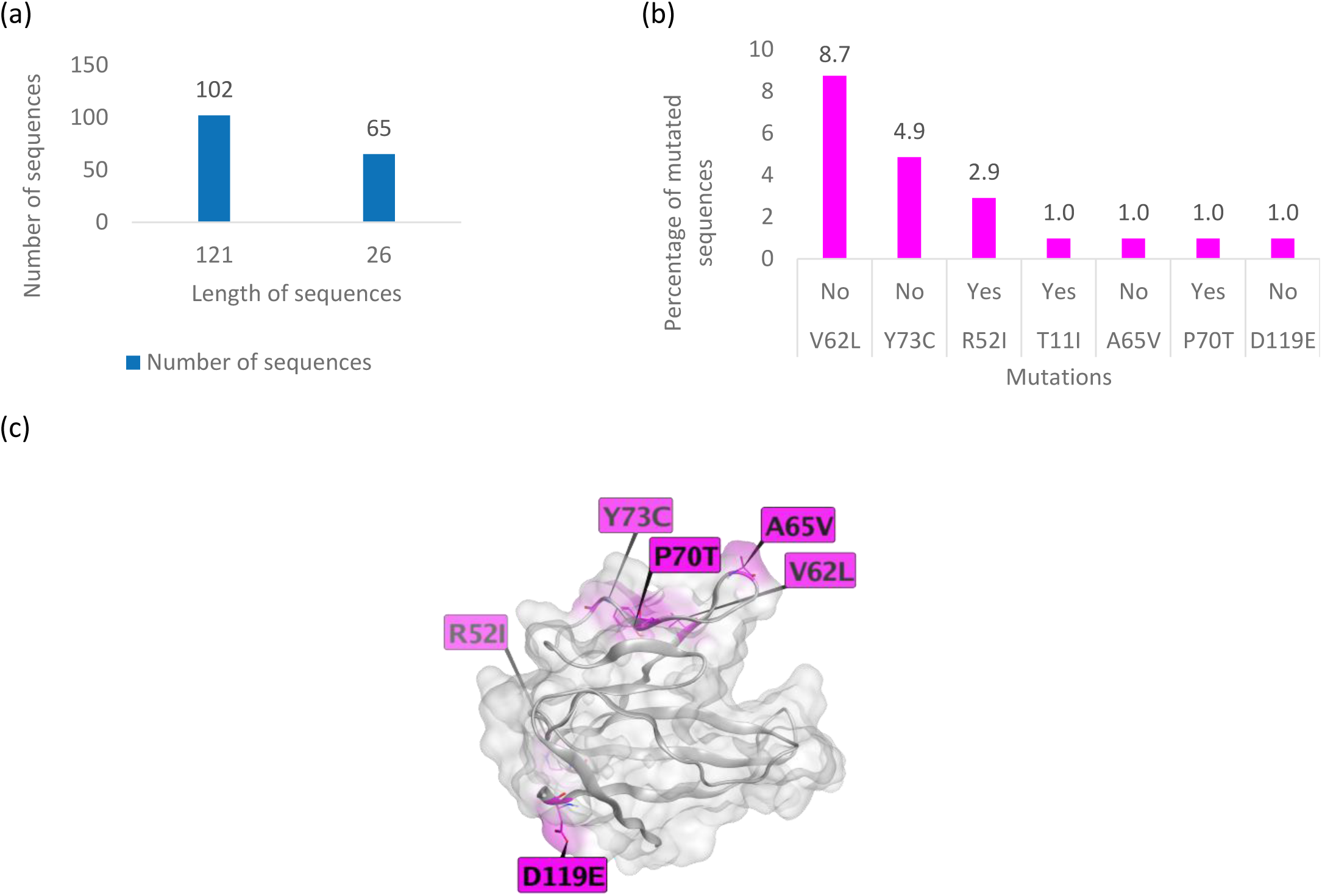
(a) Length of amino acid sequences in the Alpha variant dataset. (b) Top mutations in the Alpha variant dataset (102 sequences) showing the percentage of sequences having the mutations. Yes indicates a change in amino acid properties, No indicates no change in amino acid properties. (c) Top mutations highlighted in pink on ORF8 X-ray crystal structure 7JTL, T11I is not shown as this position is not resolved in the crystal structure.

As shown in Figure 4, the V62L, Y73C and R52I mutations, which occurred in >1 sequence, were maintained across all the variants but with low frequencies. The T11I mutation was significantly higher in the non-VOC dataset, but interestingly, it seems to have first occurred in the Alpha variant and then emerged as a dominant variant in the non-VOCs.

**Figure 4.**
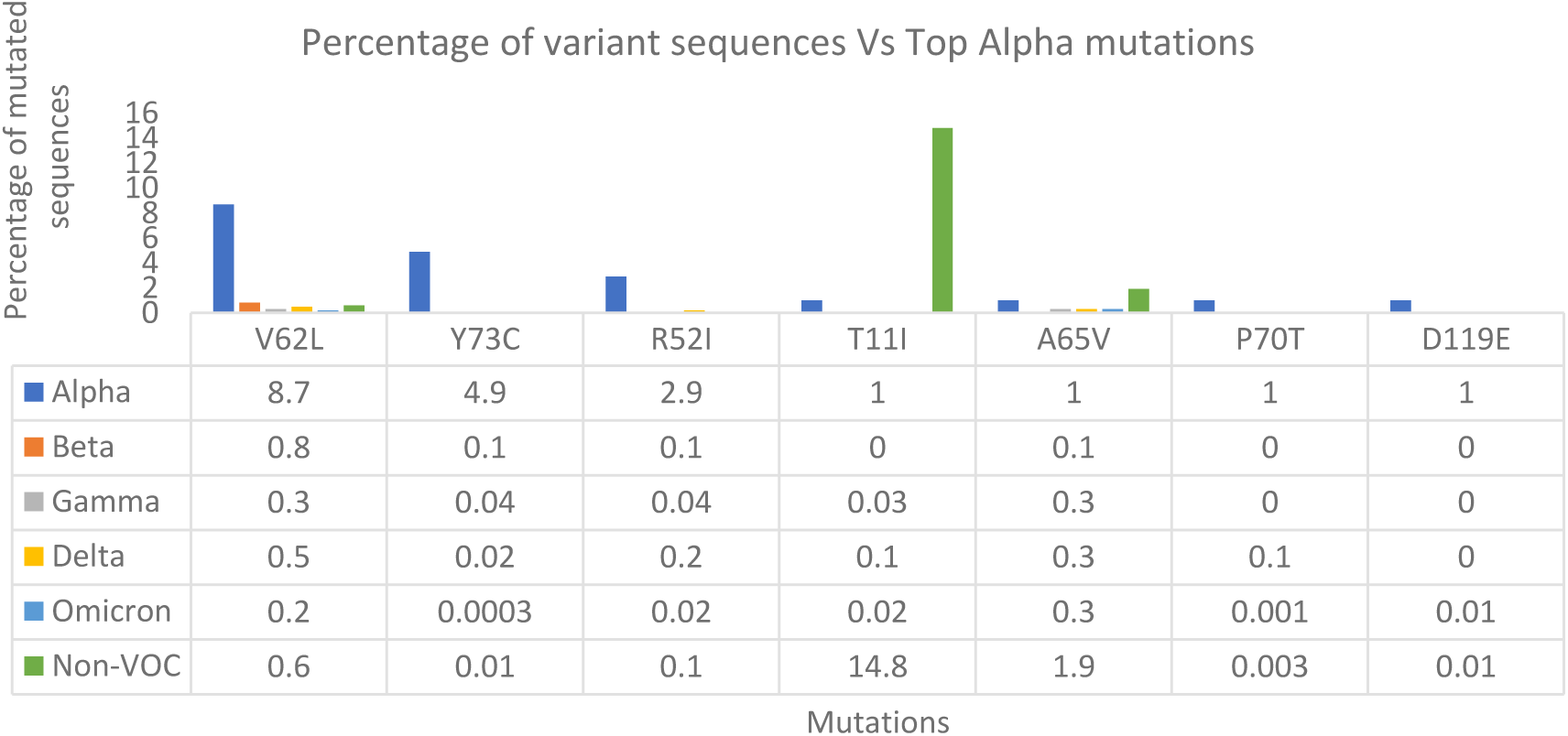
Top Alpha mutations across other variant datasets.

**(b) Beta**

The Beta variant dataset had a total of 707 sequences of length 121 AAs and 2 sequences of both 126 and 119 AA lengths (Figure 5a). The MSA was performed on all the sequences (711) and the top 10 mutations found were I121L, V62L, R115L, S67F, V81I, F120L, I121F, A51V, T87I and P93H (Figure 5b, 5c). The percentage of mutated sequences across the top 10 mutations is very low considering the small size of the Beta variant dataset e.g., the last three mutations only represent 2 sequences each. I-Mutant scores suggested that the mutations decreased the stability of the protein structure.

**Figure 5.**
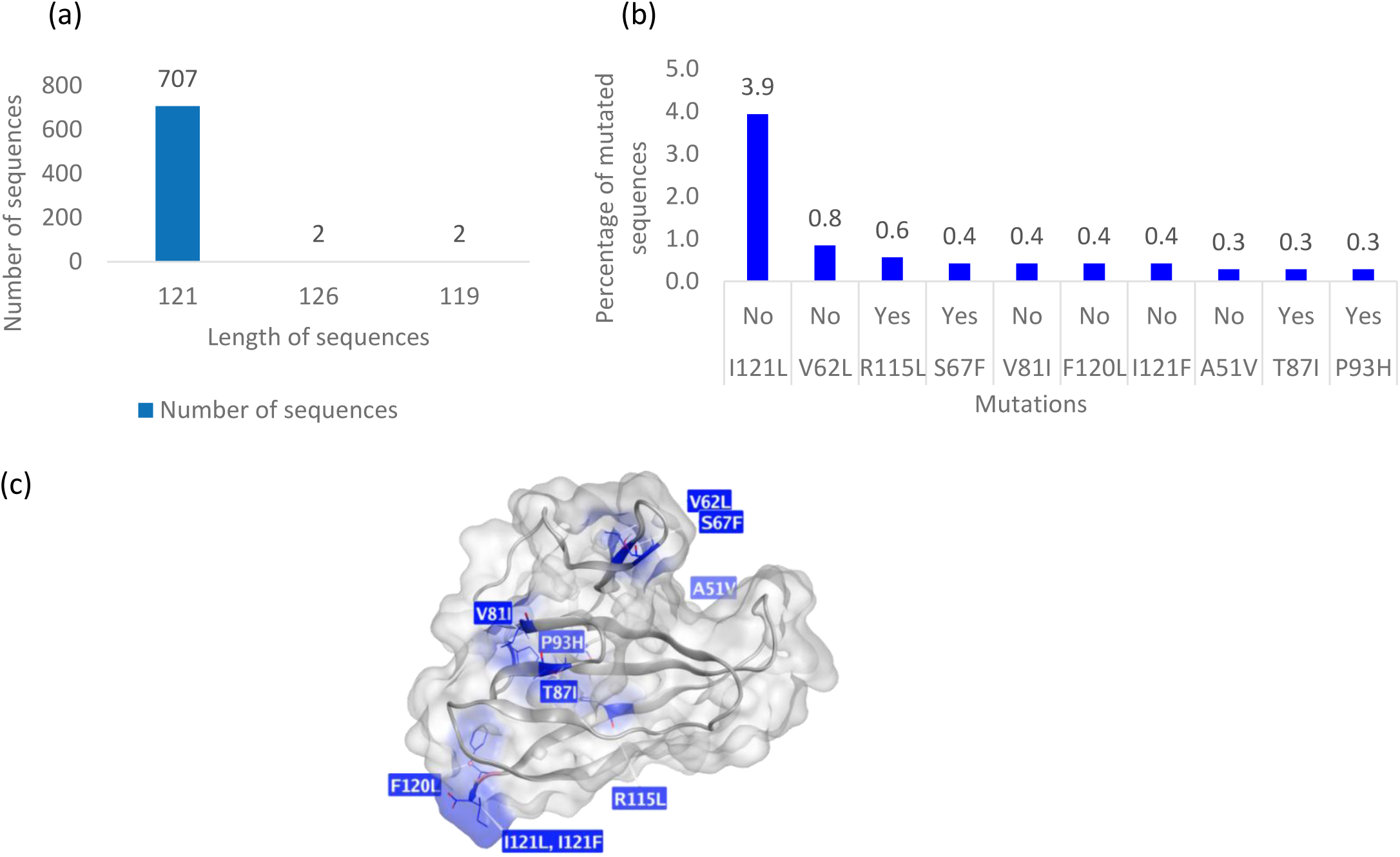
(a) Length of amino acid sequences in the Beta variant dataset. (b) Top mutations in the Beta variant dataset (711 sequences) showing the percentage of sequences having the mutations. Yes indicates a change in amino acid properties, No indicates no change in amino acid properties. (c) Top 10 mutations highlighted in blue on ORF8 X-ray crystal structure 7JTL.

As illustrated in Figure 6, for the top 10 Beta variant mutations, only the V62L mutation was maintained across the variants. The top Beta mutation I121L was present in low frequency in other variants whereas the S67F mutation frequency was observed to increase in the following variants.

**Figure 6.**
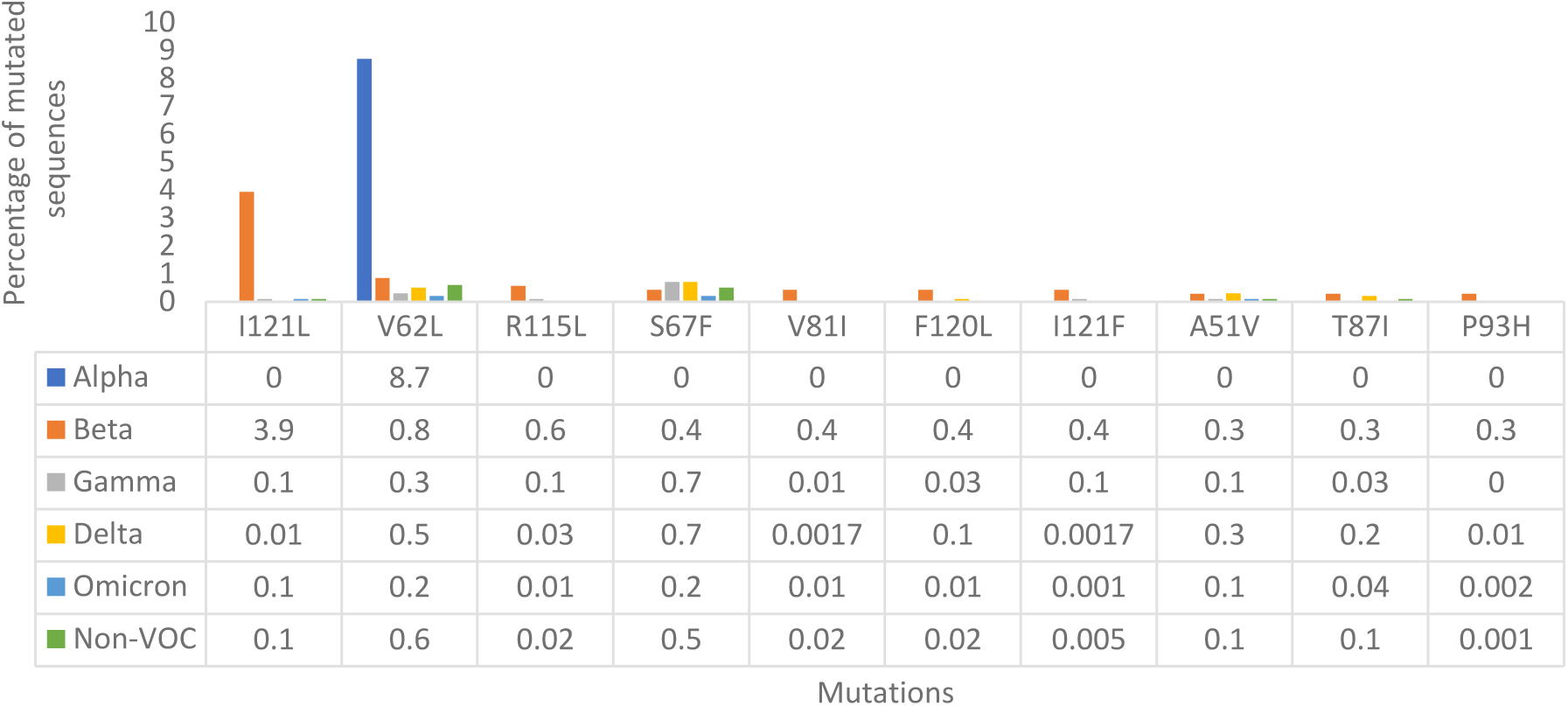
Top Beta mutations across other variant datasets.

**(c) Gamma**

The Gamma variant dataset had a total of 13,508 sequences of length 121 AAs followed by a tiny fraction of sequences with varied AA lengths (Figure 7a). The MSA was performed on all the sequences (13,535) and the top 10 mutations found were E92K, S67F, A65S, R101H, A65V, V62L, A51T, Q72H, S54L and F120V (Figure 7b, 7c). The last three mutations, Q72H, S54L, F120V were found in 28, 27 and 21 sequences, respectively. I-Mutant scores suggested that the mutations decreased the stability of the protein structure.

**Figure 7.**
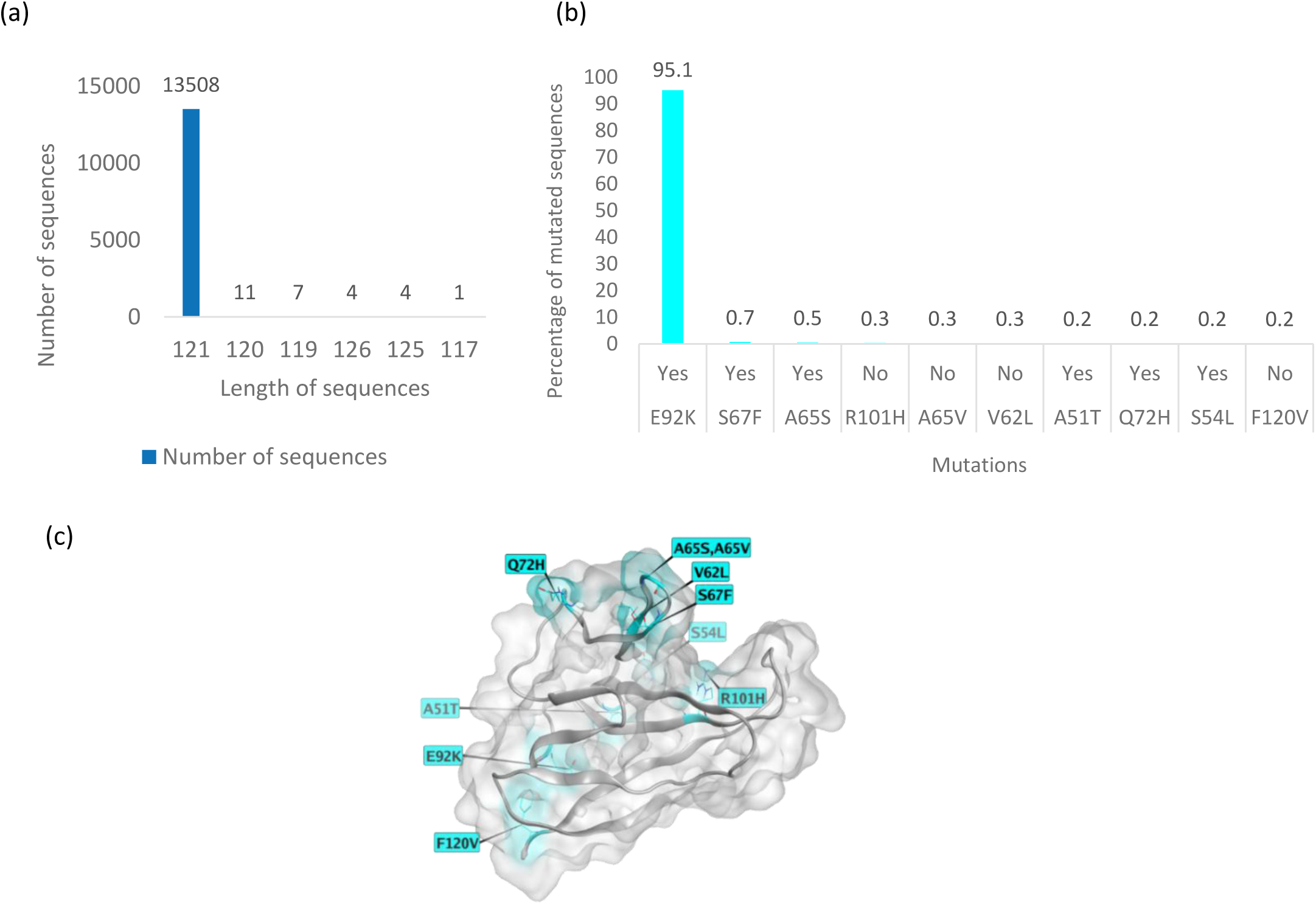
(a) Length of amino acid sequences in the Gamma variant dataset. (b) Top mutations in the Gamma variant dataset (13,535 sequences) showing the percentage of sequences having the mutations. Yes indicates a change in amino acid properties, No indicates no change in amino acid properties. (c) Top 10 mutations highlighted in cyan on ORF8 X-ray crystal structure 7JTL.

The A65V and V62L mutations were maintained at a low level across the variants. The top Gamma mutation E92K, was present in 95% of the Gamma sequences, and is very rarely observed in other variants, so it can be considered to be a Gamma signature (Figure 8).

**Figure 8.**
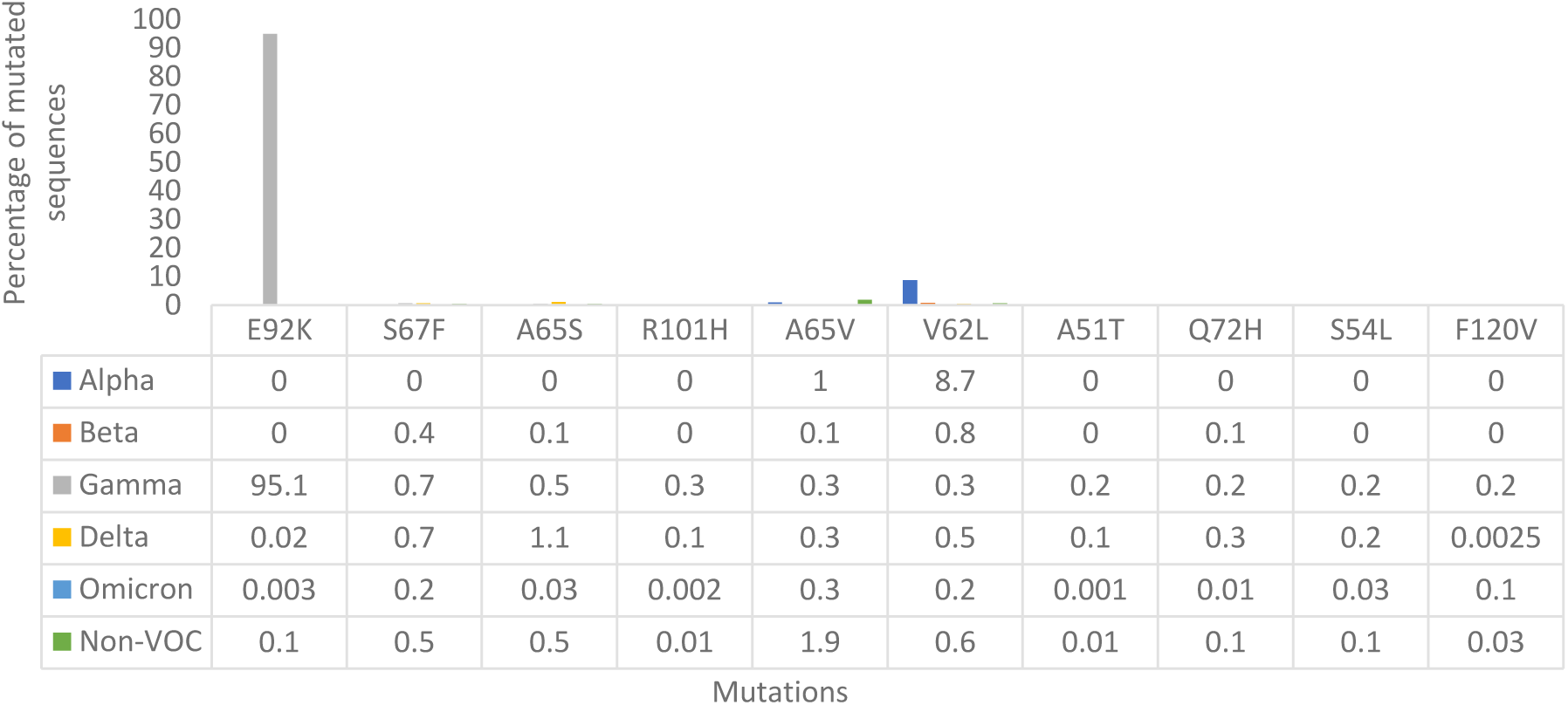
Top 10 Gamma mutations across other variant datasets.

**(d) Delta**

The Delta variant dataset had a total of 232,577 sequences of length 119 AAs followed by 7,731 sequences of length 121 AAs (Figure 9a). The MSA was performed on all the sequences (241,486) and the top 10 mutations/deletions found were F120-, D119-, L60F, Y31F, C25Y, P36S, R115C, A65S, S67S, and V62L (Figure 9b, 9c). I-Mutant scores suggested that the mutations decreased the stability of the protein structure.

**Figure 9.**
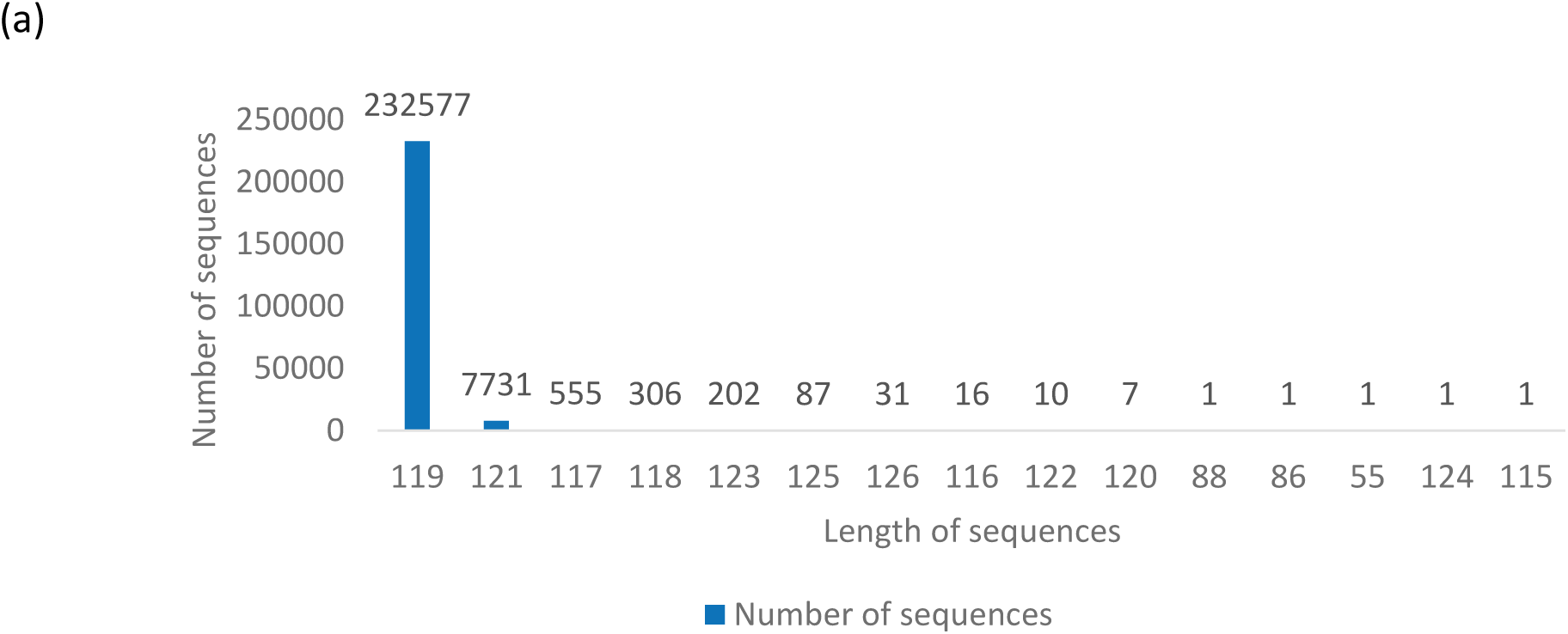
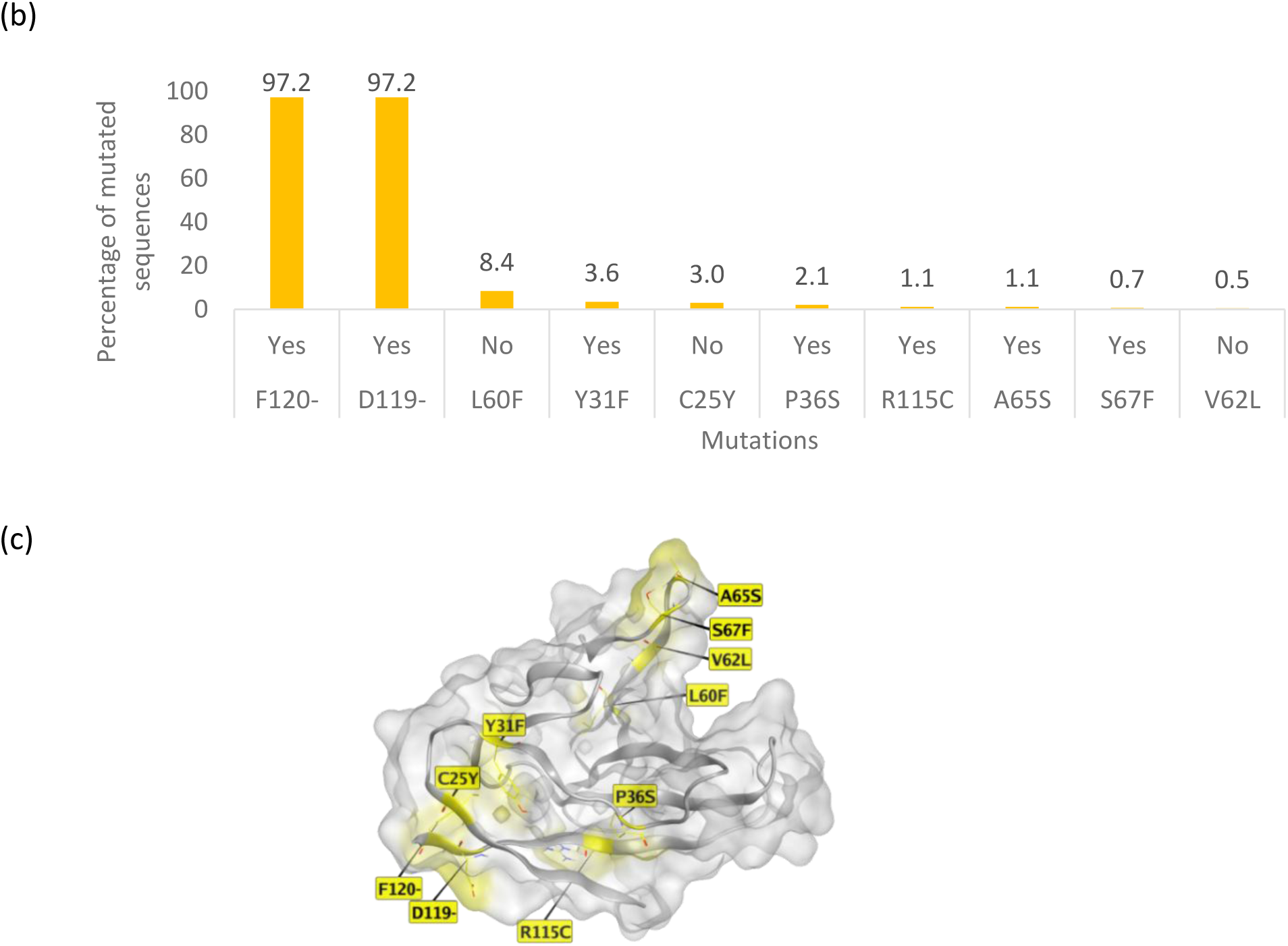
(a) Length of amino acid sequences in the Delta variant dataset. (b) Top mutations in the Delta variant dataset (241,486 sequences) showing the percentage of sequences having the mutations. Yes indicates a change in amino acid properties, No indicates no change in amino acid properties. (c) Top 10 mutations or deletions highlighted in yellow on ORF8 X-ray crystal structure 7JTL.

**Figure 10.**
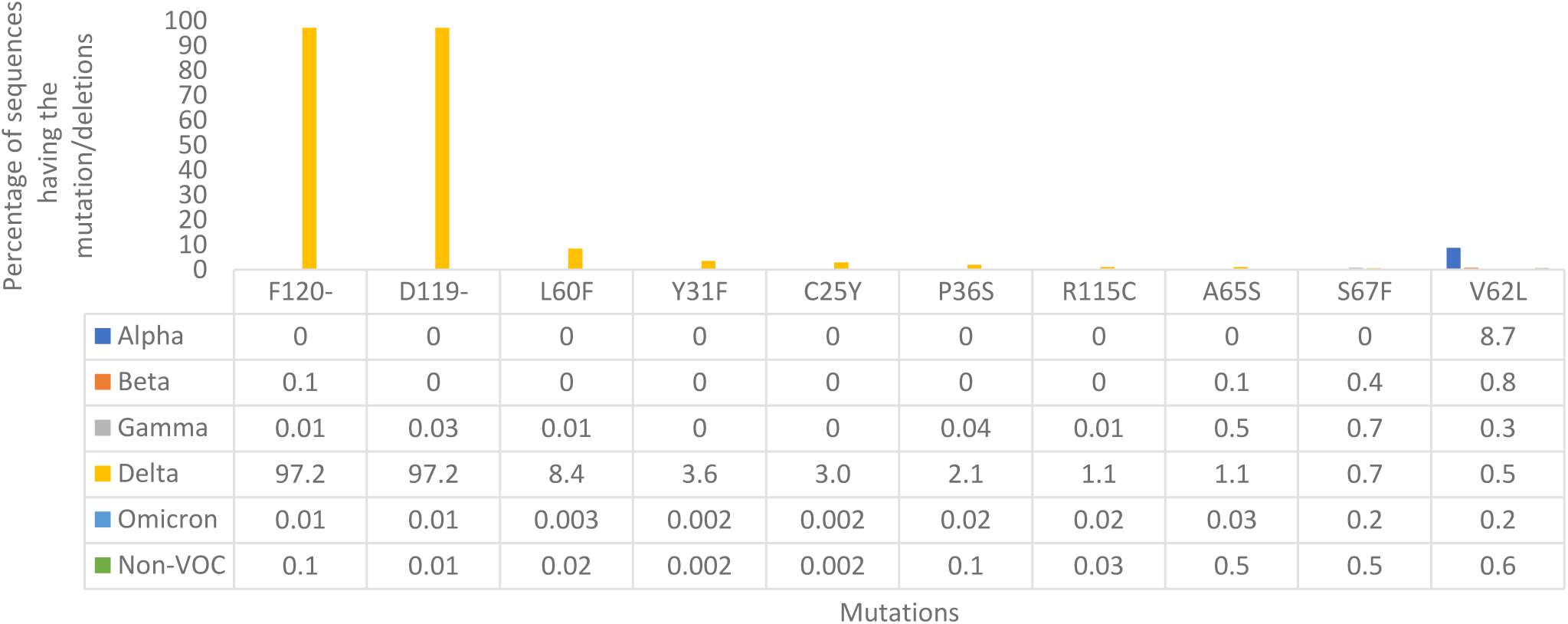
Top Delta mutations/deletions across other variant datasets.

Only the V62L mutation was maintained across the variants. The two deletions D119-and F120-were present in >97% of the Delta sequences, so they could represent a signature deletion of this variant.

**(e) Omicron**

The Omicron variant dataset had a total of 579,822 sequences of length 121 AAs followed by 394 sequences of length 119 AAs (Figure 11a). The MSA was performed on all the sequences (580,324) and the top mutations were A65D, A65V, V62L, S67F, I121L, F120V, F41S, E106D, P38S and K68E (Figure 11b, 11c). I-Mutant scores suggested that the mutations decreased the stability of the protein structure except for K68E mutation, whichincreased the stability of the protein structure. To the best of our knowledge, the biological or functional significance of this has not been reported.

**Figure 11.**
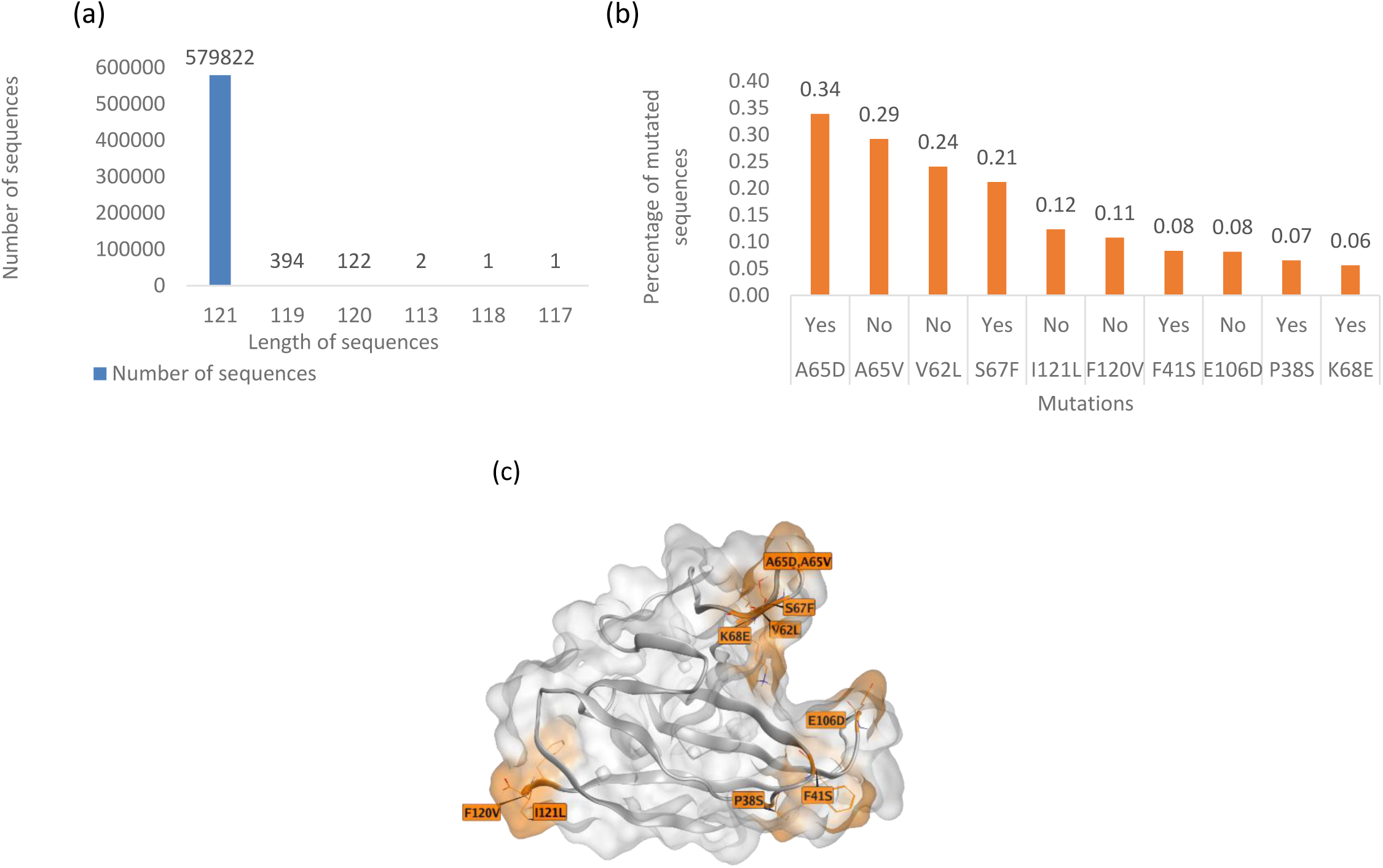
(a) Length of amino acid sequences in the Omicron variant dataset. (b) Top mutations in the Omicron variant dataset (580,324 sequences) show the percentage of sequences having the mutations. Yes indicates a change in amino acid properties, No indicates no change in amino acid properties. (c) Top 10 mutations highlighted in orange on ORF8 X-ray crystal structure 7JTL.

**Figure 12.**
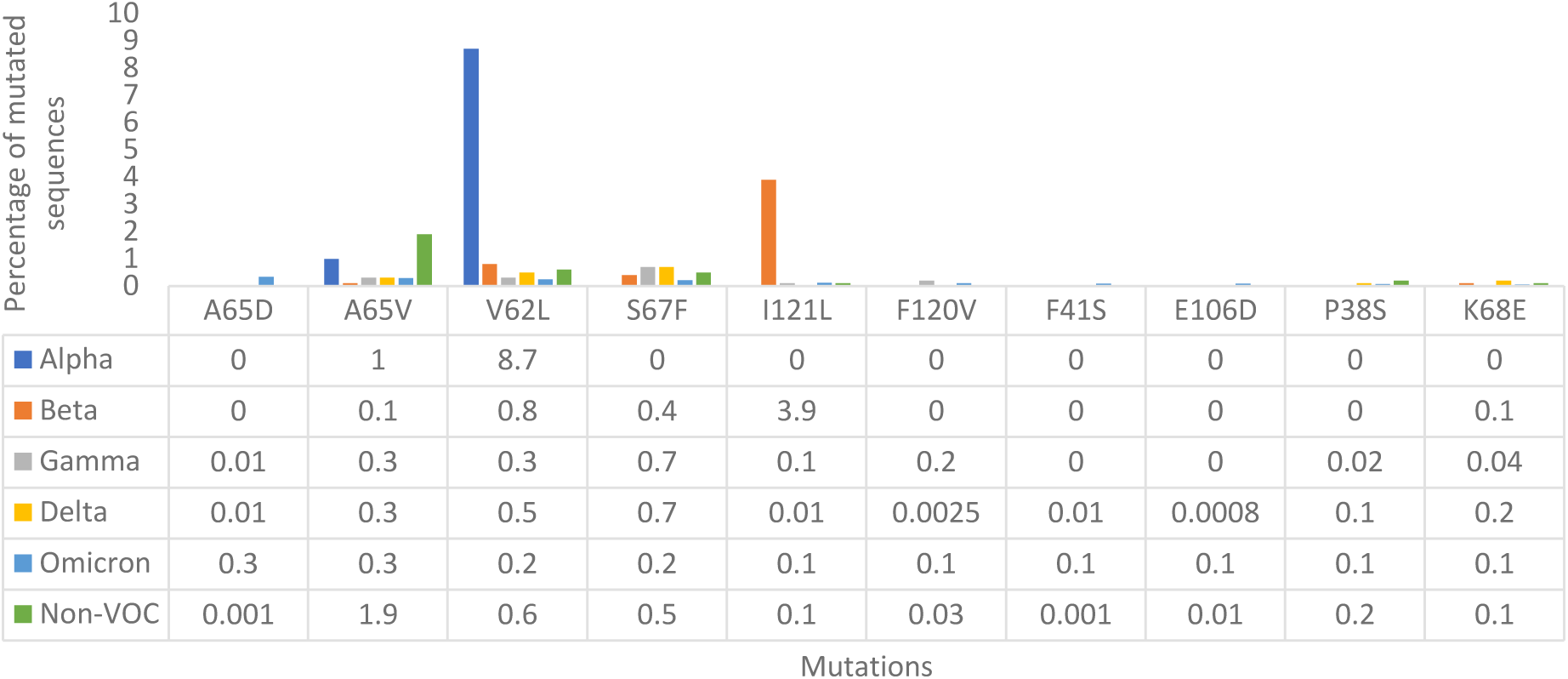
Top Omicron mutations across other variant datasets.

The A65V and V62L mutations were maintained across the variants. The top Omicron dataset mutation A65D was also present in Gamma, Delta and non-VOC datasets but with lower frequency. The relatively low frequencies of the top mutations in the Omicron dataset suggest possible sequence or functional conservation of ORF8.

**(f) Non-VOC**

The non-VOC dataset had a total of 171,818 sequences of length 121 AAs followed by 1,161 sequences of length 126 AAs (Figure 13a). The MSA was performed on all the sequences (174,475) and the top 10 mutations found were S24L, T11I, A51S, V100L, L84S, A65V, I121S, V32L, L4P and L118V (Figure 13b, 13c). I-Mutant scores suggested that the mutations decreased the stability of the protein structure except for S24L mutation, which increased the stability of the protein structure.

**Figure 13.**
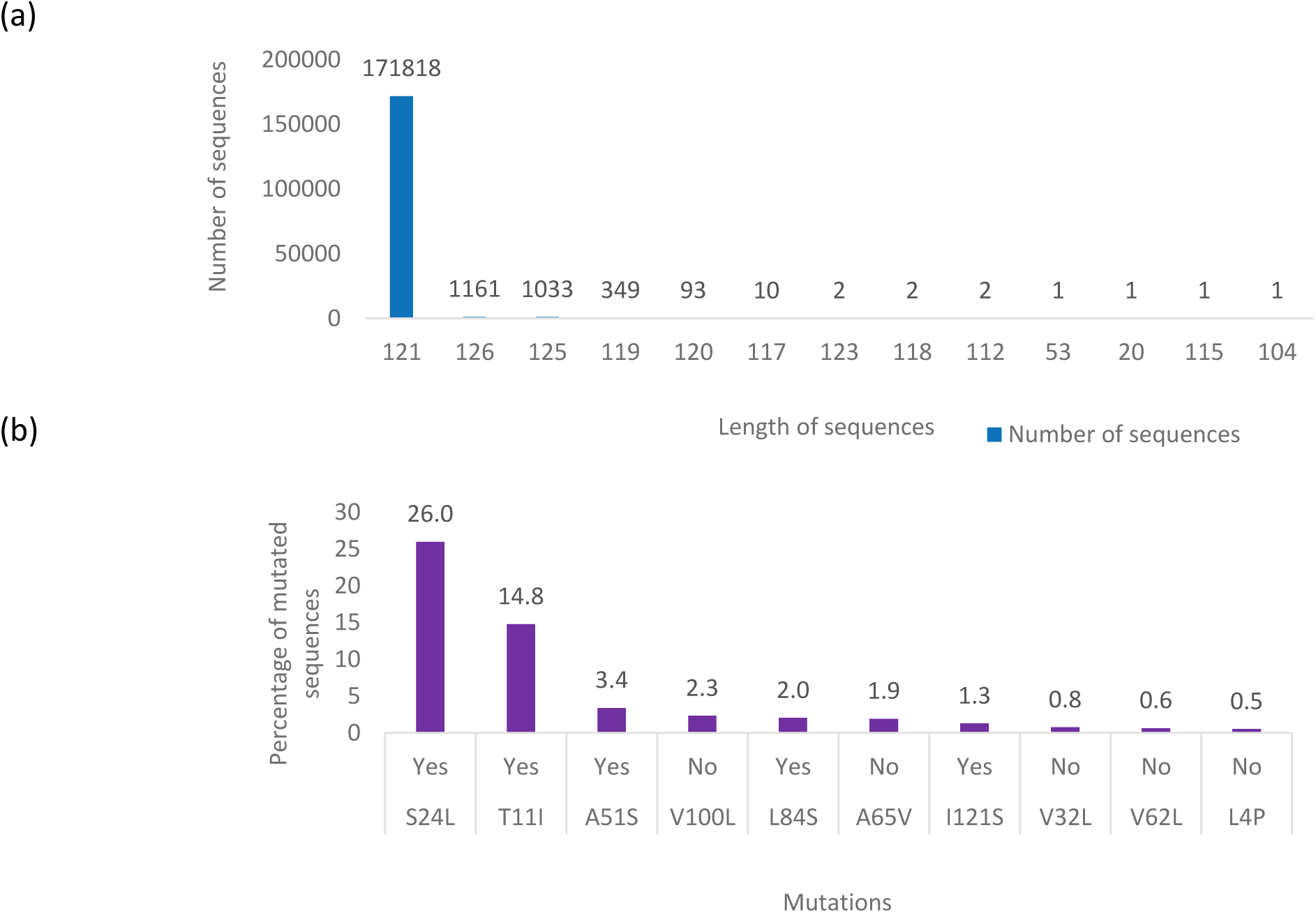
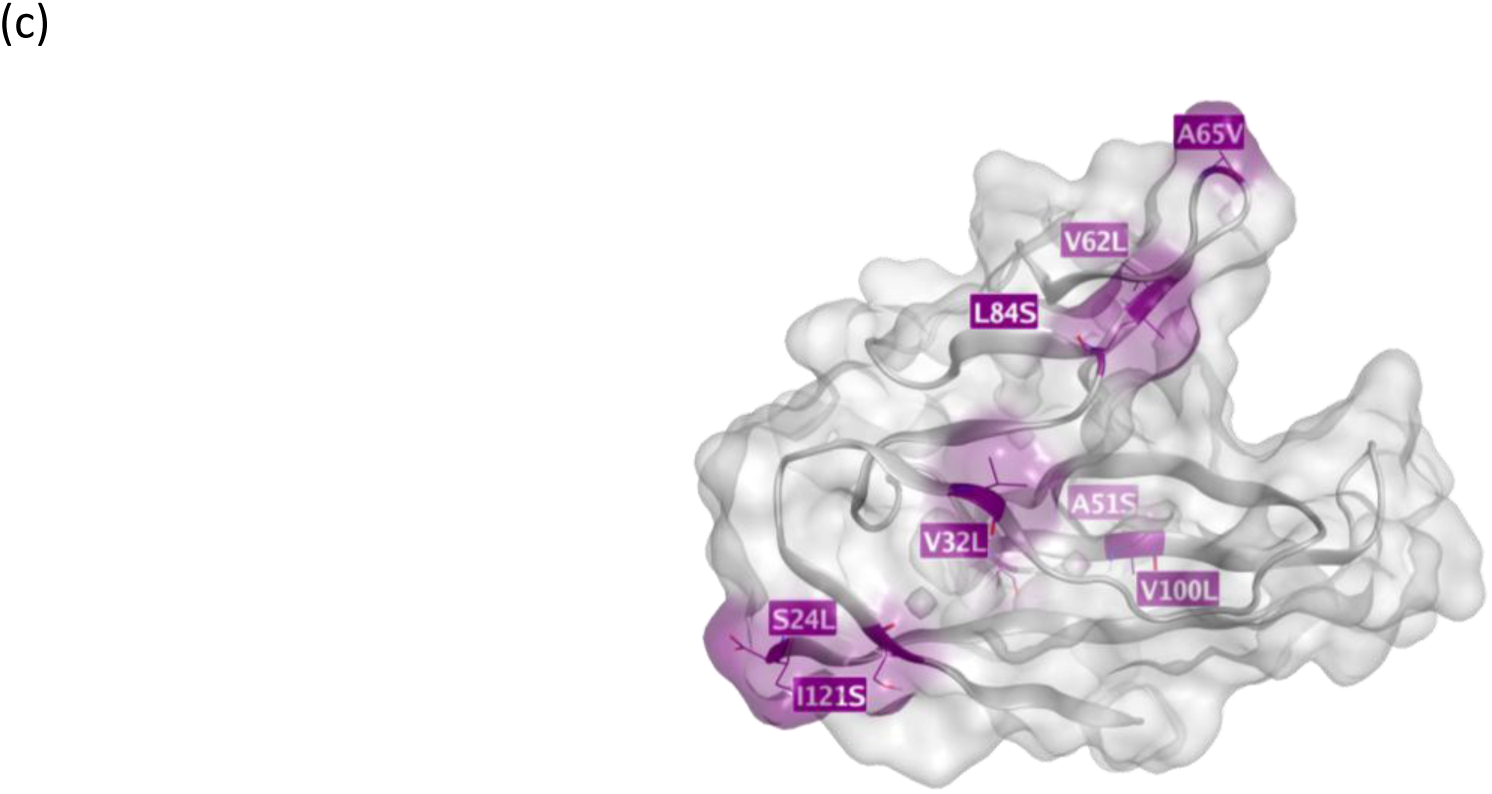
(a) Length of amino acid sequences in the Non-VOC variant dataset. (b) Top mutations in the Non-VOC variant dataset (174,475 sequences) showing the percentage of sequences having the mutations. Yes indicates a change in amino acid properties, No indicates no change in amino acid properties. (c) Top 10 mutations highlighted in purple on ORF8 X-ray crystal structure 7JTL.

**Figure 14.**
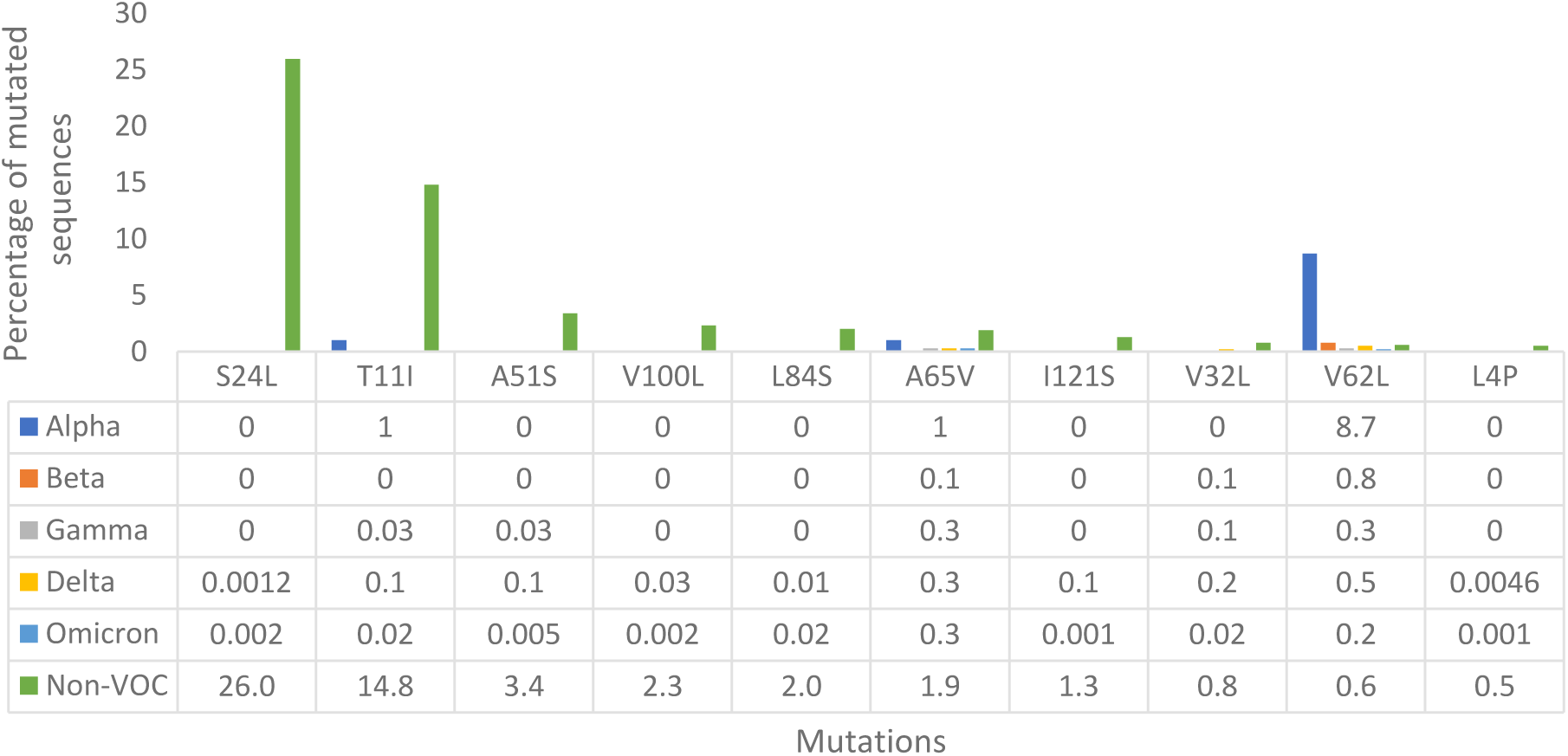
Top Non-VOC mutations across other variant datasets.

The V62L mutation was maintained across the variants and was observed to occur alongside L84S similarly to a study reported by (Laha et al., 2020). The top non-VOC dataset mutation S24L was also present in Delta and Omicron datasets but with low frequencies. In the ORF8 (1M-varying AA) sequence alignment dataset, the S24L mutation was also among the top mutations, indicating the importance of the mutation and highlighting the need for further investigation, especially as it is predicted to stabilise the protein structure.

### 2.3 Identification of conserved amino acid residues from the MSA analysis of ORF8 isolates

The MSA of eight different ORF8 isolates (Figure 15) SC-2, SC-1, mouse, bat (2 different isolates), pangolin (2 different isolates) and hypothetical (bat SARS-like) coronaviruses, showed that three amino acids (M1, L4, C102) were conserved across all the sequences. These isolate sequences were selected as they were previously reported to have high sequence identity with SC-2 Orf8 (Mohammad et al., 2020). In the ORF8 (1M-varying AA) alignment dataset, the three conserved amino acid positions had high conservation scores of 99.99%, 99.88% and 99.91%, respectively. A few motifs were also identified to be conserved, but with slight amino acid changes in ORF8 isolate sequences. Table 2 shows the top conserved amino acids from the ORF8 isolates dataset compared with the ORF8 (1M-varying AA) sequence dataset, showing corresponding amino acids that fall under conserved, intermediate and mutated clusters. From Table 2 it can be seen that M1, L7, T80, K94 and V116 were not only conserved across the ORF8 isolate sequences but also across the ORF8 (1M-varying AA) alignment dataset. Whereas the amino acid residues L4, C102, P30, V32, A55, P85 and P93 were conserved in the ORF8 isolate sequences but were found to have intermediate variability in the ORF8 (1M-varying AA) alignment dataset. Lastly, P36 was the only conserved amino acid from the ORF8 isolate sequences that was found to be mutated in the ORF8 (1M-varying AA) alignment dataset.

**Figure 15.**
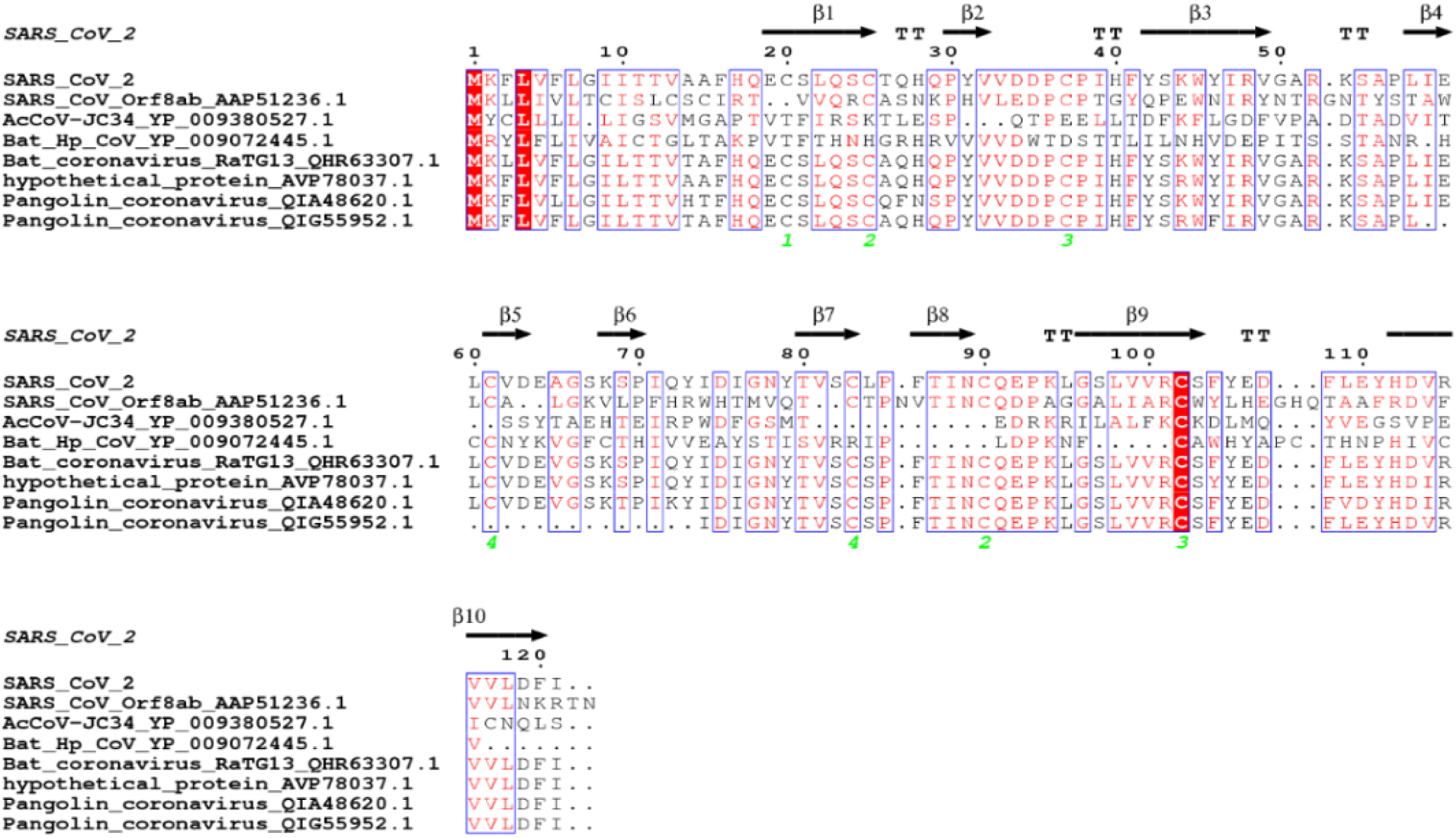
MSA of eight different ORF8 isolates showing conserved residues (red) and conserved motifs (blue box with red amino acids). The picture was generated using ESPript 3.0 (Robert & Gouet, 2014).

**Table 2.** Top conserved amino acids in ORF8 isolates dataset conservation score comparison to ORF8 (1M-varying AA) sequence dataset, along with the type of variability at the residue position and the corresponding number of sequences having that mutation represented in brackets.

| Top conserved Amino acid positions from the MSA of ORF8 isolates dataset | Conservation percent in Orf8 isolates dataset (%) | Residue position variability in Orf8 isolates dataset | Conservation/variability percent in ORF8 (1M-varying AA) alignment dataset (%) | Residue position variability in ORF8 (1M-varying AA) dataset |
| --- | --- | --- | --- | --- |
| M1 | 100 |  | 99.99 (conserved) | T (1) |
| L4 | 100 |  | 99.88 (intermediate) | P (927); F (220); I (8); V (2); A,M,R (1) |
| C102 | 100 |  | 99.91 (intermediate) | F (701); S (74); R (29); Y (25); W (1) |
| L7 | 87.5 | I (1) | 99.99 (conserved) | S (45); F (9); I (4); V (2) |
| P30 | 87.5 | V (1) | 99.92 (intermediate) | Q (260); L (230); S (69); R (62); T (52); A (1) |
| V32 | 87.5 | Deletion (1) | 99.78 (intermediate) | L (1863); A (177); I (42) |
| P36 | 87.5 | T (1) | 99.42 (mutated) | S (5214); L (520); Q (21); T (9); R (1) |
| A55 | 87.5 |  | 99.91 (intermediate) | S (146); L (66); H (38); T (4); G (2); R (1) |
| T80 | 87.5 | S (1) | 99.98 (conserved) | I (95); E (5); S (3) |
| P85 | 87.5 | Deletion (1) | 99.93 (intermediate) | S (486); L (49); T (8); H (2); R (1) |
| P93 | 87.5 | R (1) | 99.87 (intermediate) | S (744); L (217); T (174); H (39) |
| K94 | 87.5 | A (1) | 99.98 (conserved) | E (17); R (14); N (6); I,Q,T (1) |
| V116 | 87.5 | I (1) | 99.98 (conserved) | F (45); A (10); I (6); L (5); S (2) |

In order to identify more ORF8 isolates, we performed a BLAST protein-protein search (Altschul et al., 1997) (Altschul et al., 2005) on the reference SC-2 ORF8 sequence (accession ID - YP_009724396.1) resulting in 80 different aligned ORF8 isolates (see SI). The top conserved amino acids across these ORF8 isolates were C90, Q91, G96, L98, R101 and D113, with a conservation score of 98.8%. When compared to the ORF8 (1M-varying AA) sequence dataset, these residues were found to be conserved except for R101, which was in the intermediate cluster. The amino acid positions listed in Table 2 were also compared, and it was found that positions P85, P93, C102 and V116 were conserved across the 80 ORF8 isolates with a conservation score of 97.5%.

### 2.4 3D structural analysis of amino acid changes at the ORF8 dimer interface

The ORF8 3D structure is a dimer linked with a Cys20-Cys20 covalent bond (Flower et al., 2021) and the protein-protein interface (Fayne, 2013) has an area of approximately 12,260.1 Å^2^. The dimer interface has mutated or deleted amino acids, including S24, E92, D119, F120 and A51, whereas amino acids such as K53, C20 and Q18 were conserved (Figure 16).

**Figure 16:**
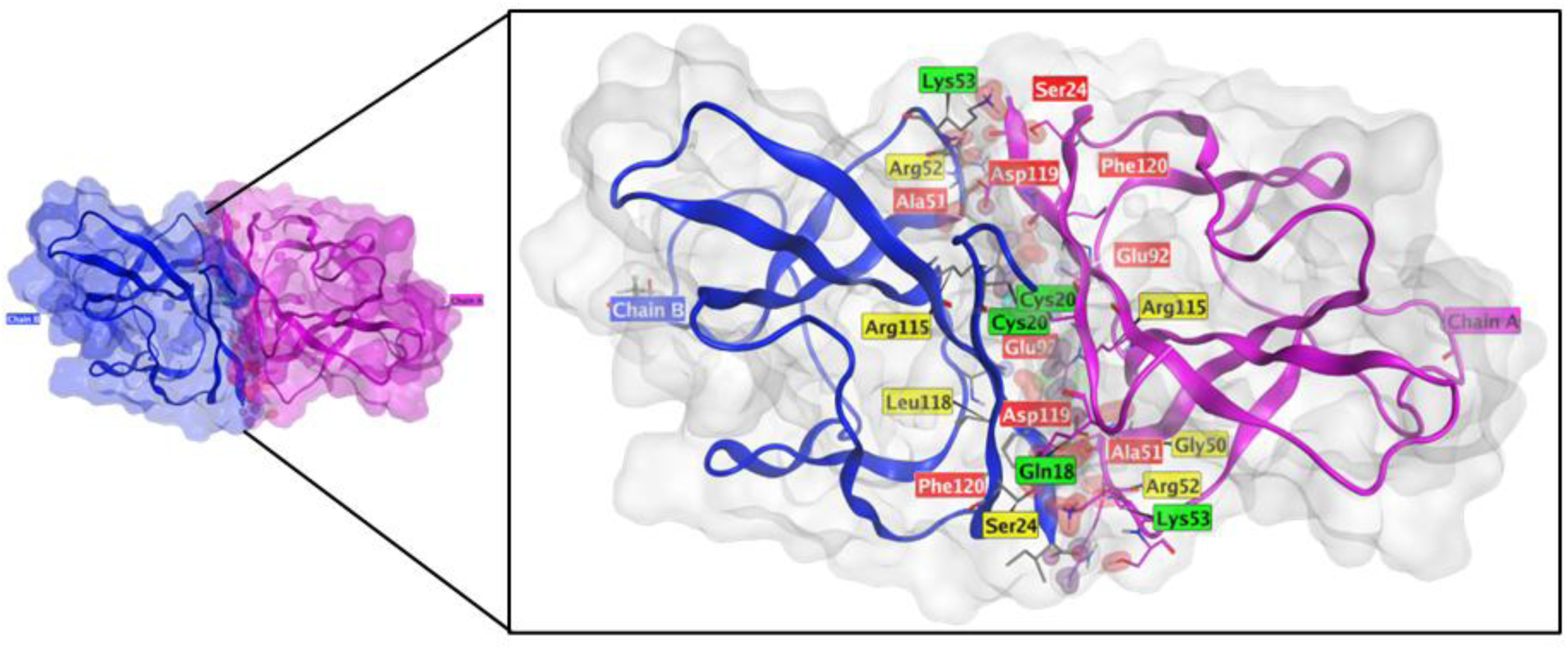
ORF8 dimer highlighting the contact residues (PDB ID: 7JTL). Purple and pink are the two ORF8 chains, red amino acids (mutated), yellow amino acids (intermediate) and green amino acids (conserved).

#### 2.4.1. Visualisation of conserved and mutated amino acid residues

From the output of the ORF8 (1M-varying AA) sequence alignment dataset, we created different clusters representing highly conserved amino acid residues, mutated amino acid residues and amino acid residues with intermediate variability as per the thresholds selected as mentioned in section 5.4. The most variable/mutated amino acids were the top 8 missense mutations and the D119/F120 deletions. On the other hand, there were 47 conserved amino acids and 64 residues with intermediate variability (Figure 17). This image nicely illustrates the structural distribution of amino acids, showing the sites that are conserved, variable, and mutated.

**Figure 17:**
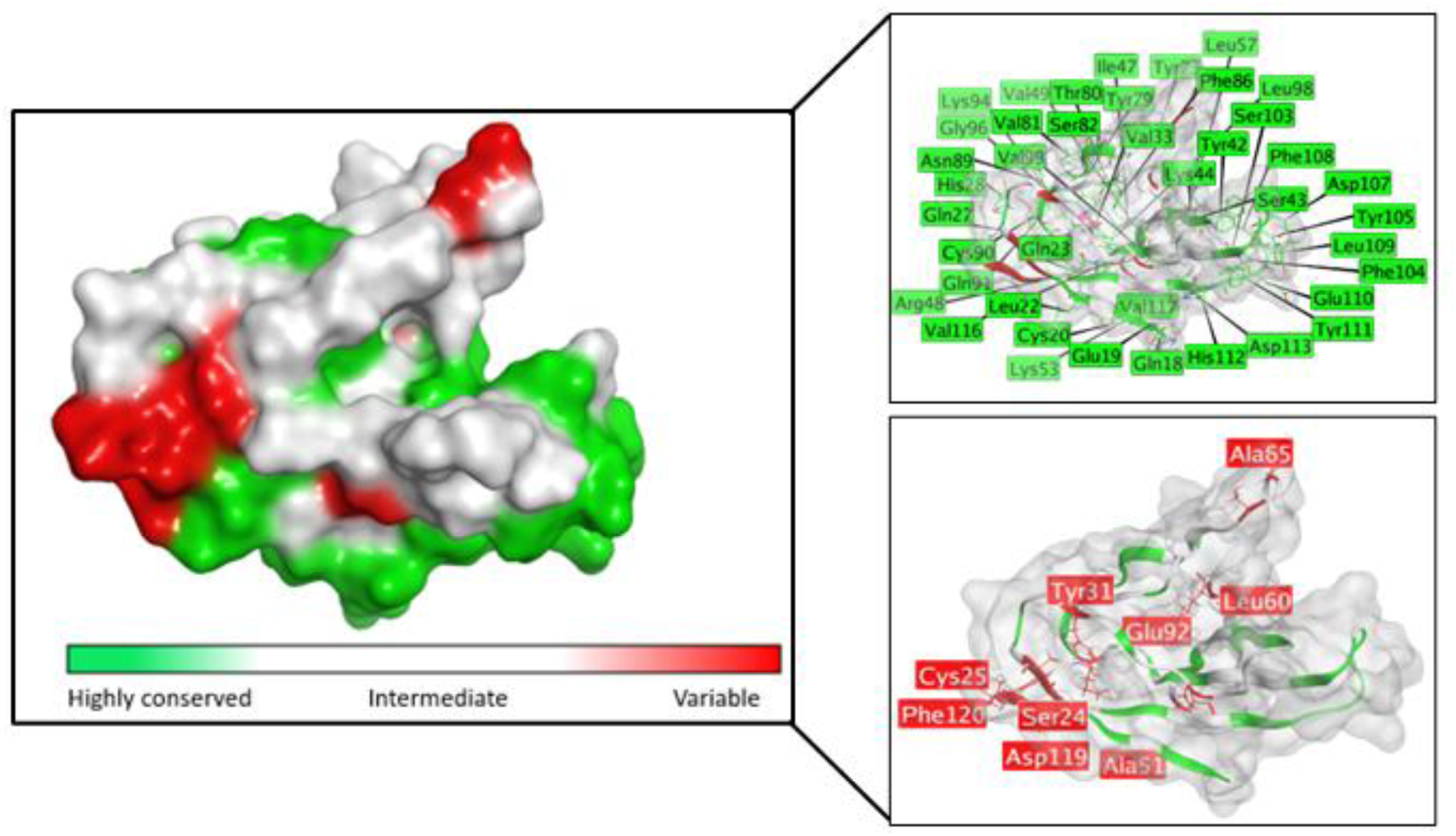
Visualisation of conserved (green), intermediate (white) and variable amino acids (red) on ORF8 X-ray crystal structure PDB ID: 7JTL. Inset: Conserved amino acid name with residue position number (green), mutated amino acid name with residue position number (red) with the exception of T11, where the position was not resolved in the PDB structure.

#### 2.4.2. Identification of conserved and mutated sites

Little is known about the binding sites present within the ORF8 protein and, mechanistically, how it interacts with human partner proteins. Our recent studies on respiratory syncytial virus (RSV) NS1 interacting with STAT1 (Efstathiou et al., 2024) and MERS-CoV nsp5 interacting with IRF3/KPNA4 (Zhang et al., 2024) highlighted possible molecular mechanisms by which human and viral proteins can interact. Considering the 3D location of conserved and mutated AAs, the possible PPIs (Fayne, 2013) sites are proposed to be siteSK, site DF1 and site DF2. These sites were selected considering the consensus localization of conserved/mutated amino acids across the VOCs and the non-VOC datasets. Two regions, i.e. siteDF1 and siteDF2, were identified where amino acid positions were found to be mutated and also had a loacalized presence across the different variant datasets (Figure 18a). The siteDF1 comprises L60, V62, A65, S67, K68, and L84, whereas the siteDF2 comprises S24, C25, D11, F120, and I121. These amino acids were often mutated across the different variants at siteDF1 and siteDF2, therefore indicating that they may be possible PPI binding sites. Alternatively, the residues S103, F104, Y105, D107, F108, L109, E110, Y111, H112 and D113 were found to be highly conserved residues concentrated at siteSK, pointing to a possible functional importance (Figure 18b).

**Figure 18.**
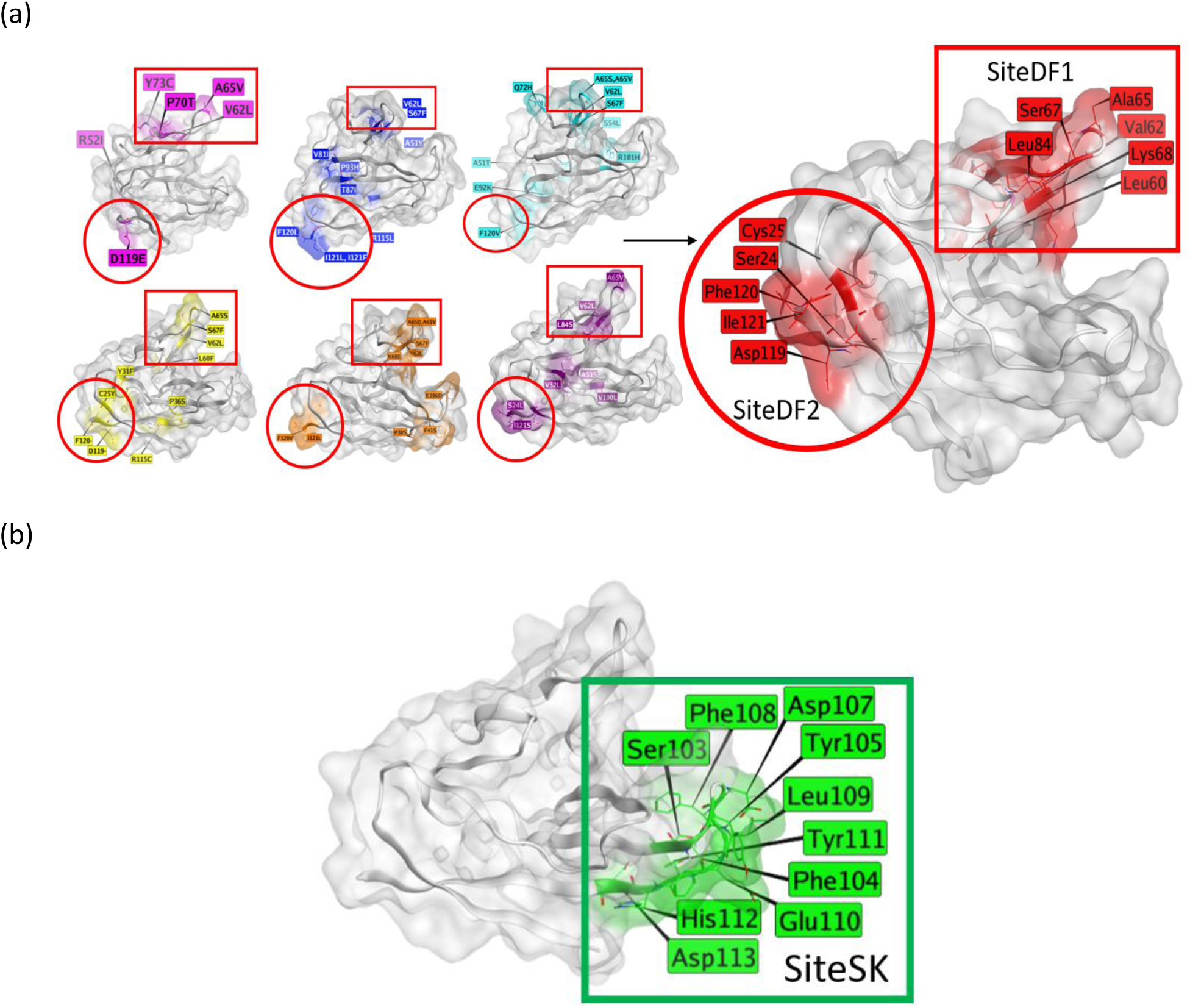
(a) Combining the main locations of ORF8 mutated amino acids across the 6 variants yields two variable regions, siteDF1 and siteDF2 (red box, red circle). Highlighted in red are the consensus mutated amino acids. (b) Conserved amino acids on ORF8 highlighting site siteSK (green box). Red: mutated residues, Green: conserved residues, Pink: Alpha, Blue: Beta, Cyan: Gamma, Yellow: Delta, Orange: Omicron and Purple: Non-VOC.

**(B) Dataset size analysis to identify most frequent mutated amino acid positions**

### 2.5 Dataset size results

In many applied scenarios, MSA is used in more of a qualitative manner to identify a set of n most-mutated positions. To gauge the reliability of such an approach on the ORF8 (700k-121 AA) dataset of 773,689 sequences, the same 100 runs of progressive resampling were conducted, quantifying how many of the top 10 mutated AA positions in the ORF8 (700k-121 AA) dataset also appeared in the top 10 within MSA of its subsets of different sizes. Based on the 100 full progressive resampling’s and MSA reruns (Section 5.6), MSA’s of the small datasets (5, 10 sequences) generally detect none or only one of the top 10 most mutated positions in the ORF8 (700k-121 AA) dataset, making them completely unreliable for finding the most mutated sites. MSAs of datasets of over 100 sequences generally detect half of the top 10 mutated positions, of 5,000 sequences detect 9 of the top 10 and of 50,000 sequences generally determine all of the top 10 mutated positions (Figure 21).

**Figure 21:**
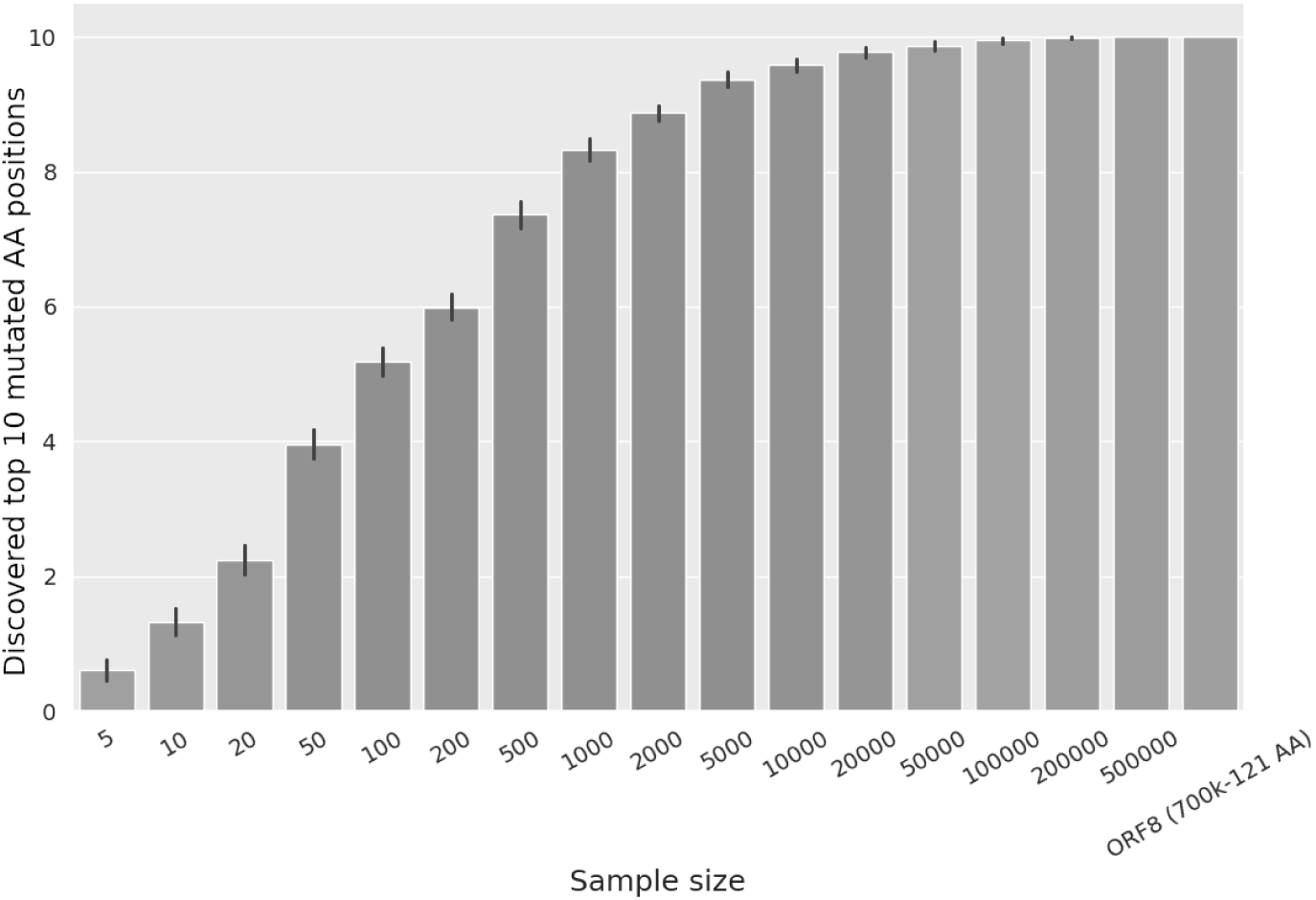
Graph showing the effect of dataset sizes (x-axis) in identifying the top 10 mutated AA positions (y-axis) of the ORF8 (700k-121 AA) dataset for 100 replicate runs. The error bars represent the confidence interval (0.95) of the RMSD for each dataset size bin.

Another, more specific line of inquiry was how many ORF8 (700k-121 AA) sequences have to be included in the MSA to reproduce the exact top 10 mutated positions of the dataset in the correct order. Only MSAs of 50,000 sequences and above were sufficient to reproduce the exact top 10 mutated positions with more than a 60% success rate, and MSAs of 200,000 sequences were needed to exceed a 90% rate of exactly reproducing the ranking of the ORF8 (700k-121 AA) dataset’s top 10 mutated positions (Figure 22). Naturally, in terms of MSA sample sizes, reproducing the exact top 10 mutated positions from the ORF8 (700k-121 AA) dataset in the exact order was a much more difficult task than simply estimating the top ‘n’ mutations and requiring more than 100x more sequences to reach comparable levels of result reliability.

**Figure 22:**
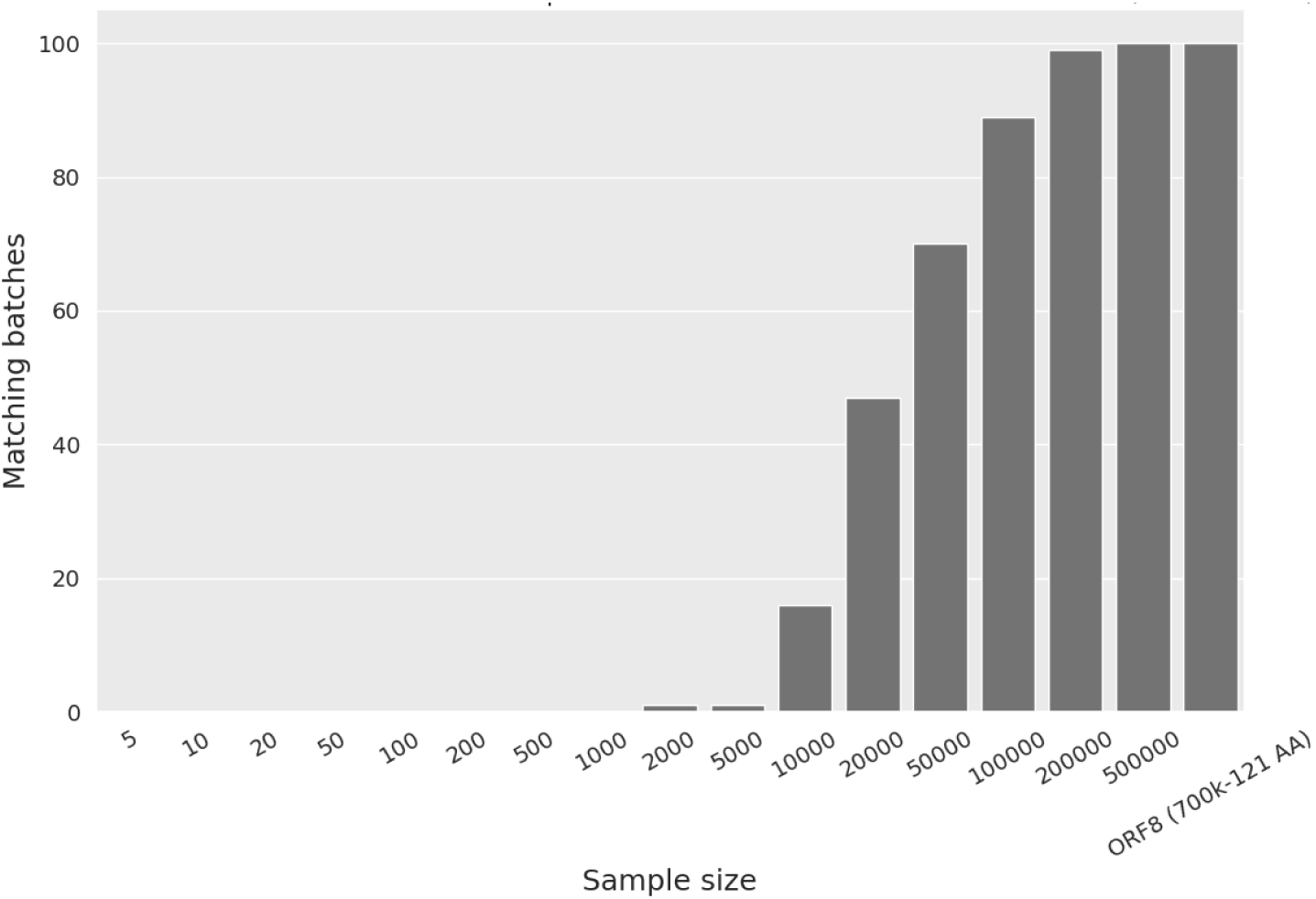
Graph showing the impact of dataset sizes (x-axis) in identifying the top 10 mutated residues from the ORF8 (700k-121 AA) dataset in the exact same order represented by the matching batches (y-axis).

Finally, a different version of the task to find the top 10 mutated AA positions was tested against the original ORF8 (1M-varying AA) dataset, which included 1,010,757 ORF8 sequences made up of all the sequences from the ORF8 (700k-121 AA) dataset and ORF8 sequences of various lengths. It was observed that none of the ORF8 (700k-121 AA) subsets were able to identify all the top 10 mutated positions of the ORF8 (1M-varying AA) dataset. The MSA of the ORF8 (700k-121 AA) dataset matched 5 out of the top 10 AA positions (Figure 23). This shows that there are 5 mutated positions i.e., S24, T11, E92, A65, and A51 that exist in the top 10 of both the ORF8 (700k-121 AA), which is gapless due to the uniform sequence size, and the ORF8 (1M-varying AA) dataset, which contains sequences of differing lengths, resulting in often large gaps in its MSA. Thus, the 5 AA positions appearing in the top 10 mutated positions in both datasets could be of particular interest, as they appear invariant to gap presence or absence in the MSAs. It is important to note here that even though the graph in Figure 23 shows 6 mutated positions, by careful examination of the MSA data it was observed that position F120, which had a deletion in the ORF8 (1M-varying AA) dataset, so due to the absence of gaps, this was identified as a missense mutation in the ORF8 (700k-121 AA) dataset. The D119-and F120-deletions were not observed in the ORF8 (700k-121 AA) dataset due to the absence of gaps. Lastly, the three remaining mutated AA positions Y31, L60 and C25 from the ORF8 (1M-varying AA) dataset were not identified within the top 10 but rather ranked low in the ORF8 (700k-121 AA) dataset analysis.

**Figure 23:**
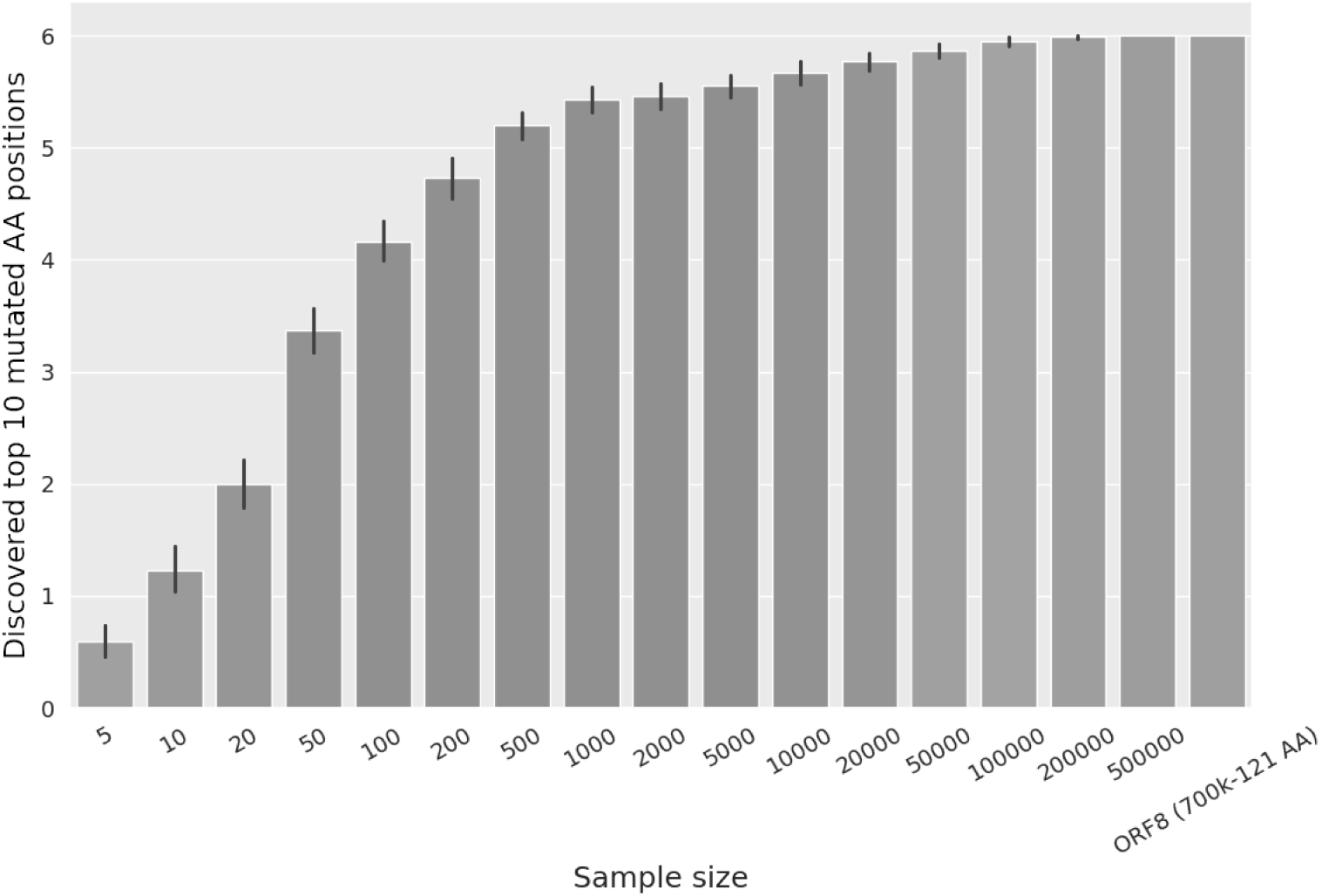
Graph showing the impact of dataset sizes (subsets of ORF8 (700k-121 AA) x-axis) in identifying the top 10 mutated AA positions (y-axis) of the ORF8 (1M-varying AA) sequence dataset. The error bars represent the confidence interval (0.95) of the count RMSD for each dataset size bin.

The rankings of the mutated AA positions are influenced by the size of the datasets and presence of varying AA length sequences, hence it is expected to observe different order of rankings when we compare the two datasets i.e., ORF8 (1M-varying AA) and ORF8 (700k-121 AA). Among the 5 common mutated amino acid positions S24, T11 and E92 have biological significance and has been discussed in Section 4.

## 3. Discussion

The ORF8 accessory protein is unique to SARS-CoV-2. Many studies have revealed its hypervariable nature (S. Chen et al., 2020)(Pereira, 2020). Datasets of small size have been used in previous MSA studies on ORF8, including a study by Badua et al., 2021, which proposed mutations from an analysis of 151 genome sequences, and another study by Liang, 2023, which also used a small dataset of 187 sequences. In contrast to these studies that identified multiple low-prevalence mutations, our study utilises a large dataset of 1,010,757 sequences, known as the ORF8 (1M-varying AA) sequence dataset, to identify mutations with high frequencies and greater statistical confidence. The multiple sequence alignment study of the ORF8 (1M-varying AA) sequence dataset revealed the presence of the top 10 mutations. These included two deletions, D119-and F120-, and eight missense mutations, S24L, T11I, L60F, E92K, Y31F, C25Y, A51S, and A65V, listed in decreasing order of mutation prevalence.

Due to their small dataset sizes, most of the studies mentioned in Table 1 overlooked the presence of L60F. However, our analysis revealed that this mutation was more prevalent and ranked 5th in the ORF8 (1M-varying AA) sequence dataset (Figure 2). We also identified deletions with a high frequency of occurrence in our dataset, which were not visible when the dataset size was small in some studies. To further highlight the importance of dataset size, the mutations Y31F, C25Y, A51S, and A65V were among the top 10 in our ORF8 (1M-varying AA) sequence dataset analysis, but they were not identified in the smaller dataset studies mentioned in Table 1. Studies by Tang et al., 2020; Laha et al., 2020; Badua et al., 2021; and Taylor et al., 2023 used small sequence datasets and found the L84S mutation to be the most prevalent. However, our ORF8 (1M-varying AA) alignment analysis ranked this mutation 16th. Liang’s 2023 study, which used a smaller dataset size, discovered the mutations R52I, Y73C, and K2Q. However, our large ORF8 (1M-varying AA) sequence dataset analysis did not identify these mutations in the top 10. This highlights the detrimental impact that small dataset sizes can have on determining correct mutational profiles, as, even though the existing studies were able to identify mutations within small dataset sizes, they have missed out on more populated mutations across the different variants.

To identify the presence of mutations across the different variants within the ORF8 (1M-varying AA) sequence dataset, we created subdatasets of sequences belonging to particular variant categories (Alpha, Beta, Gamma, Delta, Omicron, and non-VOCs). We identified previously unreported but prevalent mutations in the top 10 of each variant dataset, including P70T (Alpha), P93H (Beta), A51T (Gamma), C25Y (Delta), and F41S (Omicron). Our variant dataset analysis revealed that V62L and A65V mutations were common across all the variant datasets, suggesting a potential preference for these mutations, which could ultimately be beneficial for the SC-2 virus. From the top 10 mutations of each variant dataset, the mutations that alter the amino acid properties are R52I, T11I, P70T, R115L, S67F, T87I, P93H, E92K, A65S, A51T, Q72H, S54L, F120-, D119-, Y31F, P36S, R115C, A65D, F41S, P38S, K68E, S24L, A51S, L84S and I121S. These mutations modify the properties of amino acids, which could potentially impact the structure and interaction profile of the ORF8 protein. For instance, an MD simulation study on L84S changed the conformation of the ORF8 dimer, impacted how proteins fold, and affected the stability of the structure as a whole. It also affected the ORF8 dimer’s interfaces by lowering the number of PPIs in the dimer (Islam et al., 2023). To fully understand the structural and functional implications of many of the listed mutations, more thorough studies are still required. By analysing the variant datasets, we discovered that the Alpha variant dataset contained only 26 amino acids in 65 sequences. The truncated ORF8 was a result of the Q27STOP mutation (Liang, 2023), which in turn results in abrogating MHC-I downregulation (Moriyama et al., 2023). The top mutation of the Beta variant was I121L; a recent study showed that the I121L and E92K mutations resulted in downregulation of surface MHC-I levels (Moriyama et al., 2023). The E92K mutation was present in almost 95% of the sequences present in the Gamma dataset, again highlighting this as a signature of the Gamma variant, which was also associated with increased inflammation in the lungs (McGrath et al., 2024). Surprisingly, the mutation seems to diminish in the following variants, such as Delta and Omicron. In the case of the Delta variant dataset, only 0.02% of the sequences had the E92K mutation, whereas, in the case of the Omicron variant dataset, 0.003% of the sequences had this mutation. Earlier studies indicated that patients infected with the gamma variant had a higher likelihood of hospitalisations and ICU admissions (Martins et al., 2021) (Naveca et al., 2021). This indicates that the E92K mutation could have increased SC-2 virulence, resulting in poorer prognosis and leading to increased hospitalisations. Further work is needed to understand the mechanistic impact of this mutant on associated human PPI networks. We identified the signature deletions at D119-and F120-in the Delta variant dataset that were also somewhat visible in the Beta (F120-), Gamma, Omicron, and non-VOC datasets. Nearly 97% of the Delta variant sequences contained these deletions, which could negatively impact dimerisation abilities given the presence of D119 and F120 at the dimer interface and possibly influence virulence. To the best of our knowledge, we are the first to highlight the high prevalence of these signature deletions in a large Delta variant dataset. As previously reported, D119 and F120 play a role in dimerisation by forming hydrogen bonds and salt bridges (Flower et al., 2021); for visualisation, refer to. The Omicron VOC has a lot fewer mutations than the other variants; only 0.34% of the sequences have the most common A65D mutation (Figure 11b). This points to the possibility of the Omicron variant being less virulent. Previous reports (Wolter et al., 2022) (Vihta et al., 2023) have associated Omicron variant infections with more frequent asymptomatic carriage, milder symptoms, and decreased hospitalisation and mortality rates compared to infections with other SC-2 VOCs. In the non-VOC dataset, the S24L mutation was found in 26% of the sequences. The T11I mutation was found in almost 15% of the sequences, coming in second. The two mutations were reported by most of the studies mentioned in Table 1 but no biological consequences were discussed. These mutations might play a pivotal role in enhancing the ability of the SC- 2 virus to spread by inhibiting the MHC-I (Moriyama et al., 2023) (Wang et al., 2021). The T11I mutation was reported to down regulate surface MHC-I levels (Moriyama et al., 2023). A recent study discovered that the S24L mutation lowers the expression of a polymeric Ig receptor (pIgR), which is the same as the wild-type protein (SC-2 Orf8 S24). This shows how important this conserved activity of ORF8 is for SC-2 mucosal immune evasion (Laprise et al., 2024).

We also examined eight ORF8 isolates to identify conserved amino acids, shown in Table 2. Then, we compared the presence of these conserved amino acids to the ORF8 (1M-varying AA) alignment. The amino acids that were conserved between the two sets of data were M1, L7, T80, K94, and V116. Another analysis of the 80 ORF8 isolate sequences obtained after a BLAST protein-protein search resulted in the most conserved amino acids C90, Q91, G96, L98 and D113, which were also conserved in the ORF8 (1M-varying AA) dataset. The presence of these conserved amino acids in various isolates of ORF8 and the ORF8 (1M- varying AA) dataset, which consists of the different SC-2 variants, suggests their potential importance in influencing viral pathogenicity and immune evasion. Further research is required to elucidate their exact impact. To look at the conservation aspect of our analysis, we found 47 conserved residues with a conservation score of 99.99%-99.97% in the ORF8 (1M-varying AA) sequence dataset. This suggests that these conserved residues might have some functional importance. A subset of the conserved amino acid residues—S103, F104, Y105, D107, F108, L109, E110, Y111, H112 and D113—were found to be grouped together at one location (siteSK) (Figure 18b), which suggests that this location may be functionally important. The amino acids from this site are not present at the dimer interface and are free to interact with the host proteins, as discussed below.

Two additional potential binding sites were also identified, i.e., DF1 and DF2 (Figure 18a), considering the localisation of amino acid positions that were mutated across the SC-2 variant dataset. Interestingly, SiteDF2, comprising of residues including C25, S24, D119, F120 and I121, is present at the dimer interface, so mutations there will impact the dimerisation of ORF8. The two deletions, D119 and F120, identified in our analysis can cause disruption in the dimerisation of the two monomeric ORF8 proteins, which decreases SC- 2 viral pathogenesis. On the other hand, the SiteDF1 with the amino acid residues L60, V62, A65, S67, K68 and L84 is not present at the dimer interface but rather at a distant location, which might play a role in PPIs.

The concentration of the conserved and mutated residues in different locations suggests their potential involvement in interactions with various partner proteins. Alternatively, because siteDF1 and siteSK are next to each other, they may form a single large PPI. ORF8 is known to interfere with host immune responses, for example, downregulating the expression of MHC-I (Zhang et al., 2021), inhibiting type 1 interferon signalling pathways (Li et al., 2020) and releasing pro-inflammatory factors via IL-17 receptor (IL17-A) interaction (Lin et al., 2021) A study found that the L84 and S84 variants of SC-2 ORF8 inhibited IRF3’s nuclear translocation. It was later shown that ORF8 interacts mechanistically with HSP90B1 to cause the generation of IFN-β and IRF3 (J. Chen et al., 2022). The two amino acid variants present at the 84^th^ position are located at siteDF1. An *in-silico* study used molecular docking followed by molecular dynamics simulation and MMGBSA based binding free energy calculations to predict the binding of SC-2 Orf8 with IRF3 (Rashid, Suleman, et al., 2021). The molecular docking study revealed that SC-2 Orf8 with S24L mutation had stronger binding affinity with IRF3 and this mutation may favour immune evasion by Orf8. In our analysis this was located at siteDF2. The three sites—siteSK, siteDF1, and siteDF2—might be involved in these kinds of interactions. However, further research is necessary to understand how ORF8 interacts with host immune proteins.

Finally, we investigated the impact of sequence dataset size on the ability of MSA to identify the most prevalent mutated amino acid positions. In this study, we utilized 773,689 ORF8 sequences with a length of 121 amino acids, referred to as ORF8 (700K-121 AA), to eliminate the impact of gaps and generate estimates based on the best-case scenario. We did not use the ORF8 (1M-varying AA) dataset in this study due to its inconsistent sequence lengths, which introduced large gaps into the analysis.

The MSA analysis of progressively resampled subsets of the ORF8 (700k-121 AA) sequence dataset provides some insight into how many sequences need to be aligned to obtain a comprehensive, consistent MSA, at least for the ORF8 protein. As expected, more sequences yield more accurate MSAs with more consistency, with diminishing returns on the number of sequences analyzed. Still, there are some notable threshold values. All of the top 10 mutated amino acid positions in the ORF8 (700k-121 AA) dataset having 773,689 sequences were usually identified in its 50,000-sequence subsets (Figure 21). The overall dataset size - MSA accuracy trend appears sigmoid-like in nature, with an inflection point somewhere between 200 and 500 sequences, where it is possible to consistently identify 7 mutated amino acid positions out of 10. However, using MSA of only 5 to 10 sequences, a scenario that is not that uncommon in other studies (Chang et al., 2020; Goud et al., 2022) generally yielded only one of the ORF8 (700k-121 AA) top 10 mutated amino acid positions, or none at all. We recommend using dataset sizes of at least 500 or 1000 sequences to reliably identify 70 to 80% of the top 10 mutated amino acid positions.

While these studies were conducted using the ORF8 (700k-121 AA) sequence dataset and the mutation extent and types can be different from other proteins, we propose that the overall trends of dataset size to MSA’s accuracy would be similar across many proteins. Furthermore, alignment of ORF8 (700k-121 AA) represents a very optimistic MSA scenario (relatively small and conserved, fixed-length sequences), and thus we speculate that MSA’s of similarly sized sequence datasets for different proteins are likely to perform even worse for the same dataset size. That said, in many research scenarios, it is often necessary to work with whatever limited data is available, as aligning even just a handful of sequences could potentially yield useful results. Nevertheless, the mutations observed in such studies should then be interpreted and relied upon with due caution.

The top 10 mutated amino acid positions of ORF8 (1M-varying AA) MSA share 5 positions with the top 10 mutated amino acid positions of ORF8 (700k-121 AA) MSA. These 5 mutated amino acid positions could be of particular interest, as they appear both in the ORF8 (1M-varying AA) dataset with different-sized sequences resulting in a significant number of large gaps, and in the ORF8 (700k-121 AA) dataset of uniformly sized sequences, where gaps are not a significant factor in its MSA. Thus, these mutations on ORF8 positions (S24, T11, E92, A65 and A51) are unlikely to be an artefact of either the gap handling strategy of ORF8 (1M- varying AA) MSA, or of the same-size sequence dataset filtering that yielded the ORF8 (700k-121 AA) dataset.

## 4. Conclusions

In our study, we have performed MSA on 1,010,757 ORF8 amino acid sequences. Many previous studies on ORF8, and indeed across the wider published literature, used very small datasets upon which to base their findings. Previous studies identified mutations, but when we ran studies with much larger datasets, these mutations were found to be very uncommon, so they were not spread across the wider population. One such example is the presence of the L84S mutation, which was not identified as a top 10 mutation in our ORF8 (1M-varying AA) dataset but was reported by most of the previous studies. Much more abundant deletions and mutations were not identified in smaller datasets, whereas we found the presence of previously unknown and highly prevalent deletions and mutations such as D119-, F120- S24L, T11I, L60F, E92K, Y31F, C25Y, A51S and A65V which have much wider implications across mutational analysis studies. We are currently examining the 16 NSPs in SC-2 to determine the mutational profile of these proteins and compare them to current literature reports.

To the best of our knowledge, no other group has undertaken a comprehensive mutation analysis across ORF8 in the current VOCs and demonstrated distinct mutational signatures between the different VOCs. The major mutation that came out of the Alpha dataset analysis was V62L. There was truncation after the 27^th^ residue, i.e., Q27*(Stop), in 26 of 167 sequences. The top Alpha mutations, V62L, Y73C, R52I and A65V were also found in the other variants, demonstrating that they gave the virus a replication advantage. The major mutations from the Beta dataset analysis were I121L (3.9%) followed by V62L (0.8%). In the case of the Gamma dataset, the signature mutation was E92K (95%). The Delta dataset had two highly prevalent deletions at the C-terminal of the ORF8 protein, D119- and F120-, both with >97% prevalence and are a signature of this variant. In contrast to this, Omicron is very highly conserved across the >500,000 sequences, with the most common mutation (A65D) only having a 0.3% prevalence. Lastly, in the Non-VOC dataset, the major mutations from the analysis were S24L (26%) and T11I (14.8%).

Most importantly, the selection of thresholds to consider a range of amino acids as conserved, intermediate variability, and mutated in this study was arbitrary. This issue is prevalent across the broader MSA literature. However, this classification of amino acids is necessary to differentiate between the most prevalent mutants and those with intermediate variability, thereby aiding in the surveillance of mutants. We cannot predict the mutations that the upcoming SC-2 variants will acquire, but exploring these variable amino acid positions can assist us in monitoring their impact on ORF8.

We also hypothesised three PPI binding sites, i.e., siteSK, siteDF1 and siteDF2 on ORF8. The sites SK and DF1 possibly interact with host proteins, whereas site DF2 might play a crucial role in the dimerization of ORF8. These sites are suggested as plausible starting points for mutagenesis studies to investigate their biological significance.

Finally, we analyzed how many ORF8 sequences have to be aligned to qualitatively reproduce the MSA results of the ORF8 (700k-121 AA) dataset of all available 773,689 ORF8 sequences, each 121 amino acids long. To almost always reproduce its top 10 most prolific mutated amino acid positions correctly, 50,000 sequences were needed. Aligning 200 to 500 sequences generally yielded about 6 to 7 of the ORF8 (700k-121 AA) dataset’s top 10 mutated amino acid positions. However, aligning 5 to 10 sequences only provides one or none of the ORF8 (700k-121 AA) dataset’s top 10 mutated amino acid positions within this ORF8 sequence set, which indicates that alignments of such a limited number of sequences need to be interpreted with caution, especially since this particular ORF8 dataset consists of relatively small and conserved, fixed-length sequences - a rather optimistic scenario by any account.

By comparing the MSA of all available 1,010,757 ORF8 sequences of different lengths with the MSA of its subset of 773,689 sequences that are exactly 121 amino acids long, we identified 5 mutated positions that are shared between the top 10 most mutated positions within both datasets. These 5 positions might be of particular interest, as they are “gap-invariant” - they appear both in the ORF8 (1M-varying AA) dataset MSA with a significant number of gaps, and in the much more rigid MSA of its fixed-length sequence subset, i.e., ORF8 (700k-121 AA), where the influence of gaps is negligible.

## 5. Material and Methods

**(A) Conservation and mutational analysis**

### 5.1. Dataset collection

The SC-2 ORF8 amino acid sequences were downloaded from the NCBI (www.ncbi.nlm.nih.gov) SC-2 resources database. Relevant filters were used to extract the sequences, such as nucleotide completeness – complete, protein – ORF8, host – humans, and collection date – December 01, 2019 to July 30, 2022. The headers of the sequences had the accession ID’s and the Pangolin lineage name. After the search, 1,010,834 ORF8 protein sequences were downloaded for performing statistical analysis. The ORF8 isolates sequences were extracted from NCBI with accession IDs AAP51236.1, YP_009380527.1, YP_009072445.1, QHR63307.1, AVP78037.1, QIA48620.1, and QIG55952.1.

### 5.2. Pre-processing of dataset

The dataset was cleaned using in-house bash scripts to remove sequences containing BJXZ ambiguous characters and uncommon amino acids such as OU within the protein sequences. The protein dataset was then divided into VOCs (Alpha, Beta, Gamma, Delta and Omicron) and non-VOC datasets using in-house sets of bash commands.

### 5.3. Multiple sequence alignment

To find the most prominent mutation locations among the ORF8 (1M-varying AA) sequence dataset, MSA was performed using Kalign 3.3.2 (Lassmann & Sonnhammer, 2005) software with default gap penalty settings. We chose Kalign to conduct the MSA because it outperformed other MSA tools in terms of speed with comparable accuracy. The calculations were performed on Ubuntu 22.04 running WSL2 on a 8 core Dell Precision 3650 Tower, i9-11900 @ 2.50GHz, with an NVIDIA RTX 3060 graphics card. The ORF8 reference sequence used for the MSA was the Wuhan reference - YP_009724396 and was placed at the top of the input FASTA files of each protein dataset. For the ORF8 isolates alignment, ESPript 3.0 (Robert & Gouet, 2014) was used.

### 5.4. Analysis

The sequences CSV file was used to plot a graph between the count and length of amino acid sequences using the Matplotlib Python library. After running the MSAs, the outputs were parsed and condensed using a Python script created by our colleague Dr. Hokamp (https://github.com/khokamp/python-scripts/blob/main/msa_result_condense.py). An MS Excel sheet detailing each amino acid residue position, including gaps, percentage of conservation (number of sequences having the same amino acid at a particular residue position) and percentage of amino acid variation (number of sequences having missense mutations, insertions, or deletions at a particular residue position) was generated. MS Excel was used to perform analysis on the Hokamp Python script output Excel file. BioAider version 1.423 (Zhou et al., 2020) was also used to perform mutational analysis to identify each type of mutation along with a log file containing the number of sequences having that mutation and stating if the mutation changed the amino acid property. The graphs were plotted, showcasing the percentage of sequences having a particular mutation in each dataset using MS Excel. The conditional formatting with three colour scales was used for highlighting highly conserved (green), variably conserved (white) and highly mutated (red) amino acid residues. We selected the mutation and conservation thresholds arbitrarily from the percentage of amino acid variation. For the ORF8 (1M-varying AA) sequence dataset, we used 0.03% as the conservation threshold and 0.74% as the mutation threshold to obtain the top 10 positions for analysis.

Different clusters were created according to these thresholds, i.e., conserved cluster, intermediate cluster and mutated cluster. For the SC-2 variant datasets, the MSAs were used to identify the top mutations after processing of the Excel sheets obtained from the Hokamp Python script and BioAider software. The impact of a mutation on the protein stability was predicted by I-Mutant 2.0 (Capriotti et al., 2005), which uses a Support Vector Machine (SVM)-based method to predict the Gibbs free energy change values due to mutations. An increase or decrease in the stability of the protein structure might cause an impact on the overall function of the protein and its biological activity.

### 5.5. Molecular modelling (visualisation)

Molecular Operating Environment (MOE) version 2022.02 (MOE, 2022) was used to visualise the conserved and mutated amino acid residues on PDB: 7JTL (Flower et al., 2021) X-ray crystal structure of SARS-CoV-2 Orf8 with a resolution of 2.04 Å. The protein structure was prepared using the MOE quick prep option with the default Amber10:EHT force field, which considers explicit hydrogen atoms, tautomeric states and possible breaks in protein structure prior to conducting restrained all-atom molecular mechanics minimisation alongside electrostatics calculations. The sequence alignment tool was used to select the amino acids and label/colour them using the residue IDs.

**(B) Dataset size analysis to identify most frequent mutated amino acid positions**

### 5.6. Multiple sequence alignment and analysis

To create another MSA with reduced gap influence, a subset of the aforementioned ORF8 (1M-varying AA) sequence dataset was compiled, containing only sequences that are exactly 121 amino acids long. This subset of fixed-length sequences, named ORF8 (700k-121 AA), contains 773,689 sequences and was then processed the same way as the ORF8 (1M-varying AA) dataset. The study’s goal was to find the exact number of sequences from the ORF8 (700k–121 AA) dataset that were needed to get the same MSA results as the ORF8 (1M–varying AA) dataset.

The sequences of the ORF8 (700k-121 AA) dataset were first randomly shuffled, then progressively split into subsets of increasing sizes by taking the first 5, 10, 20, 50, 100, 200, 500, 1000, 2000, 5000, 10000, 20000, 50000, 100000, 200000 and 500000 sequences from the shuffled ORF8 (700k-121 AA). Thus, every subset contains all the sequences of the prior, smaller sets. Each smaller set was then independently aligned using Kalign, the results of which were then compared against the reference ORF8 sequence. The comparison was conducted both quantitatively, measuring the mutation rate RMSD on individual positions, and qualitatively, comparing both sets’ top 10 most frequent mutated AA positions. To make the experiment more statistically robust, this process of ORF8 (700k-121 AA) shuffling, progressive sampling, subset MSA and analysis were repeated 100 times, for a total of 1600 subset MSA and analyses. The entire process was automated using in-house Python scripts.

## Supporting Information

S01: Orf8 nucleotide positions and corresponding amino acids

S02: MSA Excel sheet of ORF8 (1M-varying AA) dataset

S03: MSA Excel sheet of Bioaider results on ORF8 (1M-varying AA) dataset

S04: BLAST search sequence alignment of ORF8 isolates

## Acknowledgments

Irish Research Council (Research Ireland) Postgraduate and Postdoctoral Fellowships (GOIPG/2021/954 and GOIPD/2023/1294; SK and IC) are gratefully acknowledged. We thank the software vendors for their continuing support of academic research efforts, in particular the contributions of the Chemical Computing Group (CCG) and OpenEye, Cadence Molecular Sciences. The support and provisions of Dell Ireland and the Irish Centre for High-End Computing (ICHEC) are also gratefully acknowledged.

## Author Contributions

Conceptualisation, S.K. and D.F.; scientific analysis, S.K., I.C. and D.F.; methodology, S.K., I.C. and D.F.; supervision, D.F.; writing—original draft, S.K., I.C. and D.F.; writing—review and editing, S.K., I.C. and D.F.; All authors have read and agreed to the published version of the manuscript.

## Notes

### Competing Interest Statement

The authors have declared no competing interest.

